# High-efficiency base editing for Stargardt disease in mice, non-human primates, and human retina tissue

**DOI:** 10.1101/2023.04.17.535579

**Authors:** Alissa Muller, Jack Sullivan, Wibke Schwarzer, Mantian Wang, Cindy Park-Windhol, Beryll Klingler, Jane Matsell, Simon Hostettler, Patricia Galliker, Mert Duman, Yanyan Hou, Pierre Balmer, Tamás Virág, Luis Alberto Barrera, Quan Xu, Dániel Péter Magda, Ferenc Kilin, Arogya Khadka, Mathieu Quinodoz, Pascal W. Hasler, Pierre-Henri Moreau, Lyne Fellmann, Thierry Azoulay, Marco Cattaneo, Simone Picelli, Alice Grison, Cameron S. Cowan, Lucas Janeschitz-Kriegl, Ákos Kusnyerik, Magdalena Renner, Zoltán Zsolt Nagy, Arnold Szabó, Carlo Rivolta, Hendrik P.N. Scholl, David Bryson, Giuseppe Ciaramella, Botond Roska, Bence György

## Abstract

Stargardt disease is a currently untreatable, inherited neurodegenerative disease that leads to macular degeneration and blindness due to loss-of-function mutations in the *ABCA4* gene. We have designed a dual adeno-associated viral vector split-intein adenine base-editing strategy to correct the most common mutation in *ABCA4* (c.5882G>A, p.G1961E). We optimized *ABCA4* base editing in human models, including retinal organoids, iPSC-derived retinal pigment epithelial (RPE) cells, as well as adult human retinal- and RPE/choroid explants in vitro. The resulting gene therapy vectors achieved high levels of gene correction in mutation-carrying mice and in non-human primates, with an average editing of 37% of photoreceptors and 73% of RPE cells in vivo. The high editing rates in primates make way for precise and efficient gene editing in other neurodegenerative ocular diseases.

## INTRODUCTION

Age-related macular degeneration and monogenic forms of macular dystrophies cause blindness. The macula is the central part of the retina containing the fovea, which enables high-resolution color vision. Currently there is no therapy for macular degeneration that halts the degeneration of cells in the retina. Monogenic forms of macular degeneration are juvenile onset and more severe, and affected patients progressively lose their ability to read, drive or recognize faces, and become blind at the center of the visual field.

The most common monogenic form affecting 1 in 6,500 individuals is Stargardt disease^1^ (STGD) (Figure 1A), which is caused by biallelic loss-of-function mutations in the *ABCA4* gene. The ABCA4 protein is a membrane transporter localized in photoreceptors and retinal pigment epithelial (RPE) cells that prevents the accumulation of toxic retinoid compounds in the retina, which otherwise lead to the degeneration of cells in Stargardt disease^2–4^. DNA base editors, which include adenine base editors^5^ (ABEs) and cytosine base editors^6^ (CBEs), are precision gene-editing tools that allow targeted repair of single nucleotide mutations without inducing double-stranded DNA breaks. The ABE complex consists of a Cas9 nickase (nCas9) fused to a tRNA adenosine deaminase (TadA) (Figure 1B) and is guided to the target DNA site by a short guide RNA (gRNA). ABEs convert A:T base pairs to G:C base pairs in the DNA through deamination in a few base-pair window. The most common STGD-associated mutation, affecting 15% of patients^7^, is a G to A point mutation in *ABCA4* (c.5882G>A, p.G1961E) that is a potential target for ABEs (Figure 1B).

**Figure 1:**
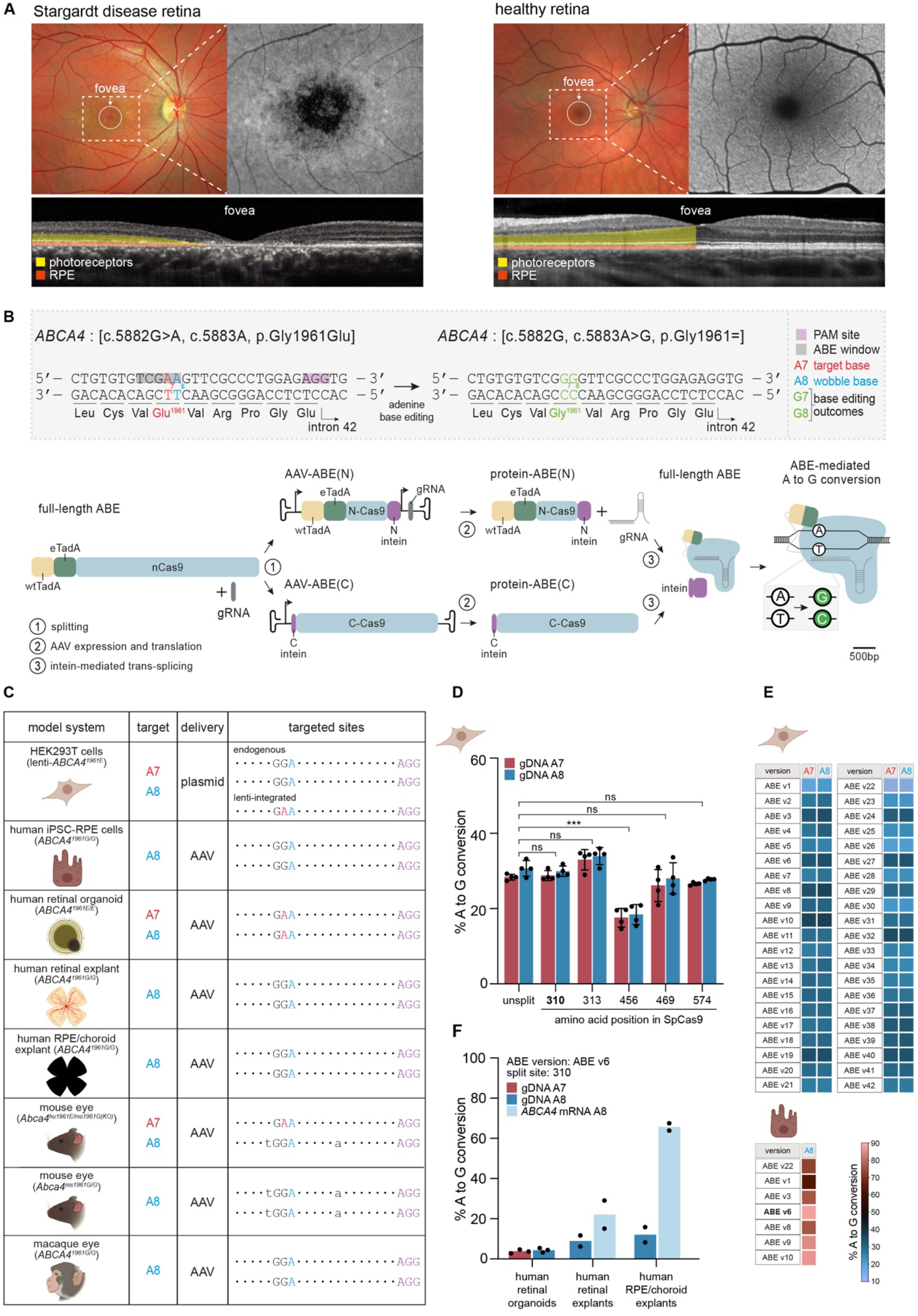
Adenine base editing corrects the most common Stargardt disease-associated mutation in vitro. **A:** Images of a retina of a Stargardt patient with biallelic *ABCA4* mutations [c.5882G>A, p.Gly1961Glu] ; [c.66G>A, p.Lys=] (left) and of a healthy individual (right). The magnified grayscale images show the corresponding autofluorescence images. Decreased foveal autofluorescence (dark region) detected in the patient indicate atrophy of RPE cells. Optical coherence tomography (OCT) images (bottom) show a cross-sectional view of the retina. The photoreceptor- and RPE cell layers are highlighted to indicate foveal thinning. **B:** Dual AAV split-intein adenine base-editing strategy. The *ABCA4* c.5882G>A mutation in exon 42 is highlighted in red, the base-editing window is indicated in grey, and the NGG-PAM site is shown in purple. Two adenines fall in the base-editing window: the c.5882A target base (A7) and the c.5883A wobble base (A8). Conversion of the A8 results in a silent mutation. WtTadA and eTadA: wild-type and evolved tRNA adenosine deaminase, respectively. **C:** Model systems used in the study to evaluate *ABCA4* base-editing efficiency in photoreceptors and RPE cells. Genotype of the model systems (first column), target adenines (second column), delivery modalities (third column) and the targeted sites (fourth column) are indicated. **D:** Base-editing efficiencies at the A7 and A8 sites in gDNA with ABE7.10 split at five different positions in lenti-*ABCA4^1961E^* HEK293T cells. Results were obtained from four replicates and presented as mean ± SD. \*\*\**P* < 0.001, by three-way mixed-effect ANOVA with Dunnett’s correction, compared to the unsplit ABE7.10 construct. ns, not significant. **E:** Base-editing efficiencies at the A7 and A8 sites in gDNA with different ABE versions in lenti-*ABCA4^1961E^* HEK293T cells (top). Results were obtained from three replicates and presented as means. Dual AAV-mediated base-editing efficiencies at the A8 site in gDNA of human iPSC-RPE cells (bottom). Results were obtained from two replicates and presented as means. **F:** Dual AAV-mediated base-editing efficiencies with the ABE v6 base editor version split at amino acid residue 310 of SpCas9 in gDNA and in *ABCA4* mRNA, in different in vitro models. Results were obtained from two to three replicates and presented as means.

Although there are reports showing the potency of base editing in cell lines and in some cases in mice^8–13^, high levels of gene correction have yet to be demonstrated in tissues of humans and non-human primates (NHPs). Here we developed an adeno-associated virus (AAV)-based adenine base-editing strategy to correct the *ABCA4* c.5882G>A mutation in clinically relevant retinal cell types and provide proof-of-concept for high-efficiency gene correction in human retina and RPE in vitro, and in mice and NHPs in vivo.

## RESULTS

To develop an adenine base-editing strategy to correct the *ABCA4* c.5882G>A mutation, we first designed and tested three different gRNAs in combination with ABE7.10^5^ (referred to as ABE v22 on Figure 1E) in a HEK293T cell line carrying the mutation on a lentivirus insert (lenti-*ABCA4^1961E^*HEK293T) (Supplemental Figure 1). We selected gRNA-3 that corrected c.5882A to c.5882G with the highest efficiency (from here on referred to as STGD-gRNA) for further experiments. STGD-gRNA places the c.5882A target base of codon 1961 at position 7 (A7) inside the base-editing window (Figure 1B). Importantly, an adjacent adenine (A8) at the 3^rd^ position of the same codon (c.5883A, wobble base) also underwent base editing (Supplemental Figure 1). Independent of A7 target base editing, the deamination of A8 does not affect the amino acid sequence (silent change) and can be used as a surrogate assay in models that lack the *ABCA4* c.5882G>A mutation (Figure 1C). This A8 position is not conserved among species (Supplemental Figure 2) and base editing of it is not predicted to interfere with splicing (MaxEntScan^14^ score = 0, dbscSNV^15^ score = 0, SpliceAI score^16^ = 0). Throughout the study, we analyzed A7 target base editing where applicable (namely, in engineered lenti-*ABCA4^1961E^* HEK293T cells, *ABCA4^1961E/E^*human retinal organoids (Supplemental Figure 3 and Supplemental Figure 4), and *Abca4^hu1961E/ms1961G(KO)^* mice (Supplemental Figure 5)). We evaluated base editing in wild-type models that are most relevant for therapeutic translation (i.e., wild-type NHPs, postmortem human retinal^17^- and RPE/choroid explants) by A8 base editing using the respective wild-type gRNA (wt-gRNA) (Figure 1C). We read out base editing at the genomic DNA (gDNA) level to reflect gene correction in all cells, and also at the *ABCA4* mRNA level to quantify gene correction in *ABCA4*-expressing cells, in photoreceptors, and in RPE cells (Supplemental Figure 6).

To deliver ABEs into the eye, we opted for AAV vectors, given the recent clinical success with an AAV2-based gene therapy ^18, 19^. As the ABE coding region exceeds the packaging capacity of AAV, we designed a split-intein ABE system^8, 20^ by separating the ABE within the *Streptococcus pyogenes* Cas9 (SpCas9) into ABE(N) and ABE(C) halves (Figure 1B). We tested five different split sites in ABE7.10 by co-transfecting plasmids expressing each half into the lenti-*ABCA4^1961E^* HEK293T cell line (Figure 1D). Base-editing rates with four of the split-intein constructs (split site 310, 313, 469 and 574) were comparable to the unsplit base editor (Supplemental Table 1, row A). For further experiments, we selected the ABE variant that is split at amino acid residue 310 of SpCas9.

ABE8 variants are evolved base editors that have higher activity in primary cells than the original ABE7.10^21^. Therefore, we compared the activity of 40 different ABE8 variants^21^ to the original ABE7.10 (v22 on Figure 1E) in the lenti-*ABCA4^1961E^* HEK293T cell line (Figure 1E). A7 and A8 editing rates showed a high correlation (*r* = 0.868) with slightly higher A8 editing. All of the engineered ABE8 variants tested led to significantly higher editing than ABE v22 (Supplemental Table 1, row B). We retained five variants (ABE v3, ABE v6, ABE v8, ABE v9, ABE v10) from this screen and tested them further in human iPSC-derived retinal pigment epithelial (iPSC-RPE) cells using dual AAV delivery (Figure 1E). We observed the highest A8 editing (up to 88%) with ABE v6. With most of the variants, we observed low level of c.5880C to c.5880T conversion due to ABE-mediated cytosine base editing^22^ (Supplemental Figure 7). This change is silent and non-conserved (Supplemental Figure 2), and is not predicted to interfere with splicing (MaxEntScan score = 0, dbscSNV score = 0, SpliceAI score = 0). Altogether, the results of these experiments identified a split-intein ABE candidate for correction of the *ABCA4* c.5882G>A mutation, namely ABE v6, split site 310.

Next, we packaged ABE v6 into an AAV9-PHP.eB capsid and evaluated base editing in *ABCA4^1961E/E^* human retinal organoids that we developed (Supplemental Figure 3 and Supplemental Figure 4), as well as in human retinal- and RPE/choroid explants (Figure 1F). To achieve simultaneous expression in cones, rods and RPE cells, we selected the ubiquitous cytomegalovirus (CMV) promoter (Supplemental Figure 8), which we combined with the rabbit β-globin polyadenylation signal (rbGlob polyA). *ABCA4^1961E/E^* human retinal organoids or human tissue samples were incubated with an equal mixture of AAV-ABE v6(N) and AAV-ABE v6(C) (2.3 × 10^11^ v.g./organoid and 3.32 × 10^11^ v.g./human tissue). Seven weeks post-transduction, we isolated bulk- gDNA and RNA. Whilst *ABCA4* editing rates in gDNA averaged between 4% and 12% in all models, higher editing rates were found for *ABCA4* mRNA, with averages of 22% and 66% in human retinal- and RPE/choroid explants, respectively (Figure 1F). These results suggest that *ABCA4* base editing can be achieved in STGD target cells, but optimization of the vector components is necessary to maximize therapeutic benefit (Figure 2A).

**Figure 2:**
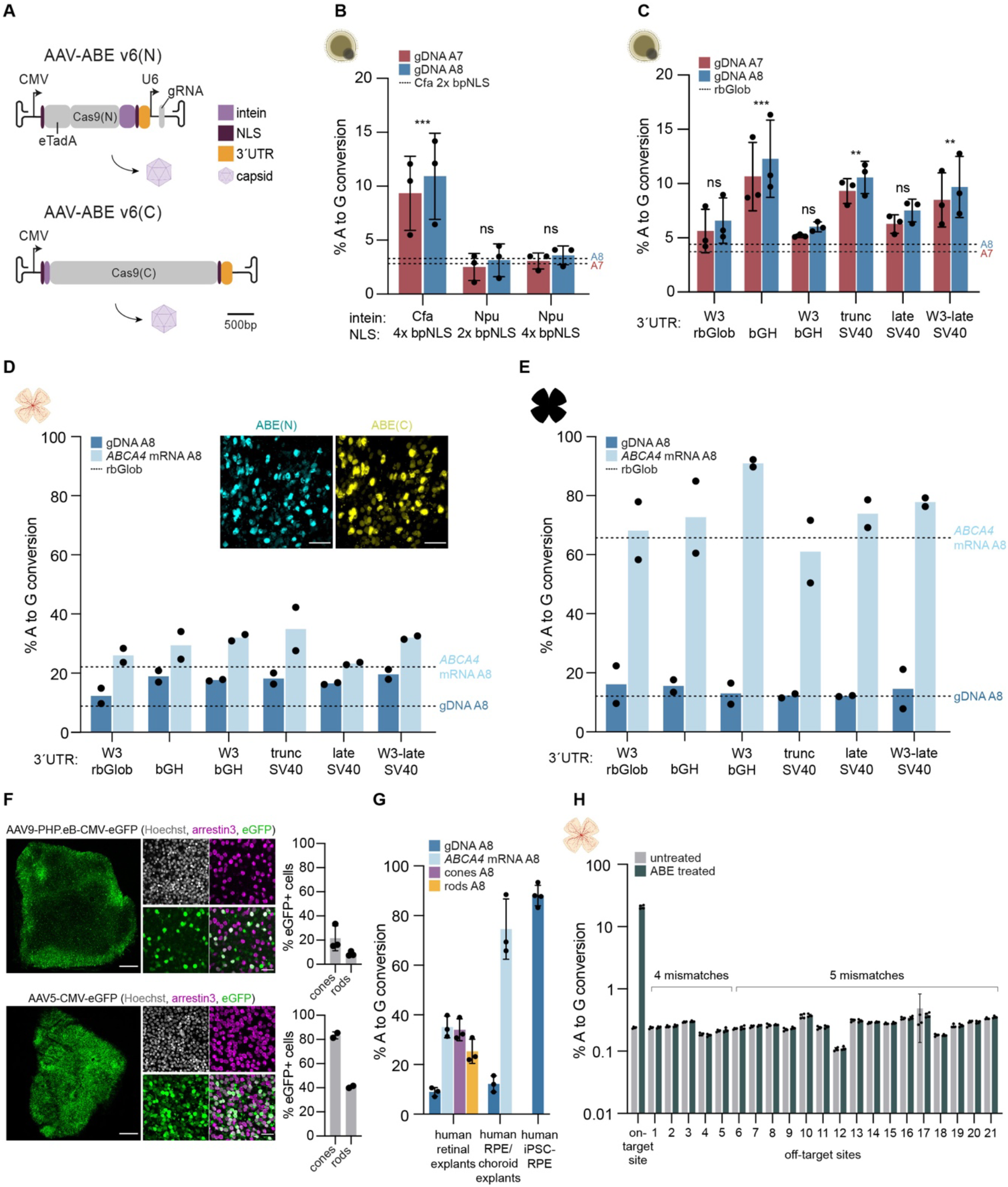
In vitro optimization of the adenine base editor AAV vectors. **A:**Schematic of the dual AAV split-intein ABE v6(N) and ABE v6(C) vectors. The variable elements in the constructs tested are highlighted in different colors. **B:** In vitro base-editing efficiencies at the A7 and A8 sites in gDNA of *ABCA4^1961E/E^*human retinal organoids with split-Cfa intein versus split-Npu intein and 2x bpNLS (one on each construct) versus 4x bpNLS (two on each construct) sequences. Results were obtained from three replicates and presented as mean ± SD. \*\*\**P* < 0.001, by three-way mixed-effect ANOVA with Dunnett’s correction, compared to split-Cfa intein and 2x bpNLS. ns, not significant. **C:** In vitro base-editing efficiencies at the A7 and A8 sites in gDNA using different 3’UTRs in *ABCA4^1961E/E^* human retinal organoids. Results were obtained from three replicates and presented as mean ± SD. \*\**P* < 0.01, \*\*\**P* < 0.001, by three-way mixed-effect ANOVA with Dunnett’s correction, compared to rabbit β-globin polyA (rbGlob). ns, not significant. **D:** In vitro base-editing efficiencies at the A8 site in gDNA and *ABCA4* mRNA using different 3’UTRs in human retinal explants and representative immunofluorescence images of ABE(N) and ABE(C) expression in the photoreceptor layer (scale bars: 25 µm). Results were obtained from two replicates and presented as means. Cyan: ABE(N); yellow: ABE(C), gamma correction has been applied to obtain optimal dynamic range for visualization. **E**: In vitro base-editing efficiencies at the A8 site in gDNA and *ABCA4* mRNA using different 3’UTRs in human RPE/choroid explants. Results were obtained from two replicates and presented as means. **F:** Representative immunofluorescence images of human retinal explants transduced with the AAV9-PHP.eB (top) or AAV5 (bottom) capsid encoding for CMV-eGFP. Left: low magnification images (scale bars: 500 µm), middle: high magnification images (scale bars: 25 µm), right: quantification of eGFP-expressing cones and rods. Values are mean ± SD. Grey: Hoechst; green: eGFP; magenta: arrestin3. **G:** Base-editing efficiencies with AAV5-SABE1 at the A8 site in gDNA and *ABCA4* mRNA of human retinal- and RPE/choroid explants, and in human iPSC-RPE cells and gDNA of sorted cones and rods. Results were obtained from three to four replicates and presented as mean ± SD. **H:** Editing rates at the *ABCA4* c.5883A on-target site and 21 predicted off-target sites in untreated human retinal explants and ABE treated human retinal explants. Results were obtained form two biological replicates sequenced twice and are represented as mean ± SD.

To optimize the vector components^23^ in vitro in human retina and RPE models, we first analyzed the effects of the type of intein and nuclear localization signal (NLS) on base-editing efficiency in *ABCA4^1961E/E^* human retinal organoids (Figure 2B). The A7 editing rates with split-consensus fast DnaE (Cfa) intein^24^ and two bipartite NLS^25^ (bpNLS) per ABE half were threefold higher than with split-Cfa intein and a single bpNLS per half. No significant difference in editing rates could be detected with split-*Nostoc punctiforme* (Npu) intein^24^ (Supplemental Table 1, row C). Next, we substituted the rbGlob polyA with different 3’ untranslated region (3’UTR) elements (Figure 2C, D and E). Immunostaining for ABE(N) and ABE(C) confirmed expression of the base editor halves in human photoreceptors (Figure 2D). The three best candidates in *ABCA4^1961E/E^* human retinal organoids and human retinal explants were bovine growth hormone (bGH) polyA, truncated simian virus 40 (trunc SV40) polyA, and truncated woodchuck hepatitis virus posttranscriptional regulatory element (WPRE) -late SV40 (W3-late SV40) polyA, which lead to a two-to threefold improvement in base editing at the gDNA level (Supplemental Table 1, row D). These candidates also performed well in human RPE/choroid explants (Figure 2E). We chose bGH polyA for further studies, as it is part of a clinically approved ocular gene therapy vector^18^, and the combination of CMV-ABE v6(N)-Cfa-(2x bpNLS)-bGH polyA-U6-gRNA and CMV-Cfa-ABE v6(C)-(2x bpNLS)-bGH polyA (denoted as Stargardt adenine base editor 1 (SABE1)) became our lead candidate ABE vector for further studies. We validated SABE1 in vitro in human iPSC-RPE cells and *ABCA4^1961E/E^* human retinal organoids in independent experiments (Supplemental Figure 9).

Next, we tested different AAV capsids to identify serotypes that efficiently transduce photoreceptors and RPE simultaneously. We packaged a CMV-eGFP construct into eight different capsids (AAV2, AAV5, AAV8, AAV8-BP2, AAV9-7m8, AAV9-PHP.B, AAV9-PHP.eB and Anc80L65) and transduced human retinal explants as well as *ABCA4^1961G/G^* human retinal organoids (Supplemental Figure 10 and Supplemental Figure 11). Major differences in photoreceptor transduction efficiency were found between different serotypes. Interestingly, in human retinal explants, AAV5 showed the highest transduction efficiency and outperformed AAV9-PHP.eB in photoreceptor transduction (Figure 2F), leading to high base-editing rates in human photoreceptors (on average 35% in *ABCA4* mRNA) (Figure 2G). We developed an intracellular immunostaining-based, fluorescence-activated cell sorting (FACS) protocol to determine editing rates in cones and rods separately (Supplemental Figure 12). Editing rates reached 34% on average in sorted cones and 25% on average in sorted rods (Figure 2G). High base-editing rates with the AAV5 capsid were also detected in human RPE/choroid explants (on average 75% in *ABCA4* mRNA) as well as in human iPSC-RPE cells (on average 88%) (Figure 2G).

To analyze genome-wide off-target effects of base editing on human tissue, we performed targeted deep-sequencing on the top 21 computationally predicted off-target sites from a base-edited human retinal explant (Figure 2H and Supplemental Table 2). Off-target sites were selected based on the number of mismatches (up to 5) between the STGD-gRNA and the GRCh38 human reference genome, as well as on the presence of a NGG-protospacer adjacent motif (PAM)^26^. Although the c.5883A (A8) on-target editing rate was on average 21% in gDNA, no off-target base editing was found for any of the 21 sites tested.

Based on the encouraging in vitro base-editor efficiency and precision in photoreceptors and RPE cells, we tested SABE1 in vivo in 11- to 22-week-old mice (Figure 3A). Stargardt disease patients with the *ABCA4* c.5882G>A mutation are almost exclusively compound heterozygous for the mutation^27^. Therefore, we created an *Abca4* mouse strain (*Abca4^hu1961E/ms1961G(KO)^*) (Figure 1C and Supplemental Figure 5) that carries a single humanized *Abca4* c.5882G>A allele (*Abca4^hu1961E^*) and an *Abca4* knock-out allele (*Abca4^ms1961G(KO)^*) in trans. Similarly to Stargardt patients, only a single *Abca4* c.5882A allele (*Abca4^hu1961E^*) can be targeted by the STGD-gRNA. Therefore, the percentage of edited cells in this model corresponds to the measured editing rates.

**Figure 3:**
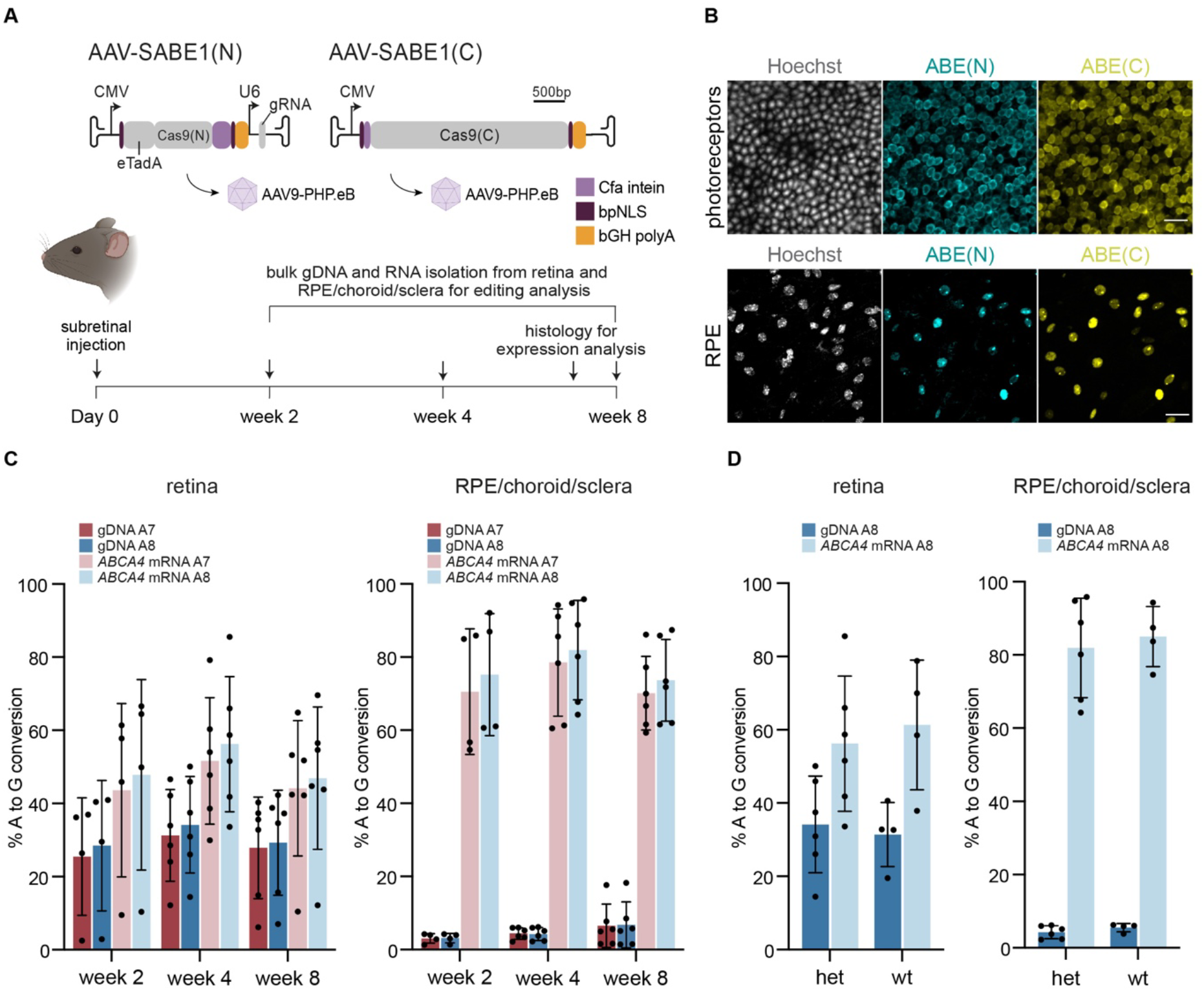
In vivo base editing in mice. **A:**Experimental design. Dual AAV9-PHP.eB-SABE1 was delivered by subretinal injection. Eyes were harvested at 2-, 4- and 8 weeks post-injection and the retina and RPE/choroid/sclera were processed separately. **B:** Representative immunofluorescence images of ABE(N) and ABE(C) expression in the photoreceptor layer (top, scale bars: 12.5 µm) and RPE layer (bottom, scale bars: 25 µm) of *Abca4^ms1961G/G^* wild-type mice at 7 weeks post-injection. Grey: Hoechst; cyan: ABE(N); yellow: ABE(C). **C:** In vivo base-editing efficiencies in *Abca4^hu1961E/ms1961G(KO)^*mice in the retina (left) and RPE/choroid/sclera (right) at different time points after treatment. Results were obtained from four to six eyes and presented as mean ± SD. **D:** In vivo base-editing efficiencies in *Abca4^hu1961E/ms1961G(KO)^*(het) and *Abca4^ms1961G/G^* (wt) mice in the retina (left) and RPE/choroid/sclera (right) at four weeks after treatment. Results were obtained from four to six eyes and presented as mean ± SD.

Given that the autofluorescent signals representing the deposition of toxic lipids was previously found to be elevated in the full *Abca4* knock-out mouse model^28^, we analyzed this phenotype in *Abca4^hu1961E/ms1961G(KO)^* mice. We found no increase in autofluorescence signals in this model compared to wild-type mice (Supplemental Figure 5).

We subretinally injected SABE1 (3 × 10^10^ v.g./eye) and analyzed expression of the two ABE halves (Figure 3B). Based on immunostaining against ABE(N) and ABE(C), we observed very high expression in both the photoreceptor layer (Figure 3B, top) and the RPE layer (Figure 3B, bottom). Most cells expressed both halves simultaneously (79% for retina and 72% for RPE) and a few cells expressed only one base editor half (1% ABE(N), 5% ABE(C) for retina and 1% ABE(N), 4% ABE(C) for RPE). We then quantified base editing from injected *Abca4^hu1961E/ms1961G(KO)^* mice in the retina and RPE/choroid/sclera tissue at 2-, 4- and 8 weeks after injection (Figure 3C and Supplemental Table 3). At four weeks post-injection, A7 editing efficiency on the single *Abca4* c.5882A allele was on average 31% for the retina and 4% for RPE/choroid/sclera gDNA. However, the editing rates in the target cells were expected to be higher because mostly photoreceptors (∼80% of all cells in the mouse retina^29^) and RPE cells (only a few percentage of all cells in the RPE/choroid/sclera) are targeted after subretinal injection. Indeed, at the *Abca4* mRNA level (which represents the relevant target cells), we observed A7 editing with averages of 52% and 79% edited transcripts in the retina and RPE/choroid/sclera, respectively. Increase in incubation time did not lead to increased base-editing rates. As previously observed in vitro, A7 editing rates correlated strongly with A8 editing rates, both in gDNA and in mRNA (Supplemental Table 1, rows E, F, G and H).

We also tested base editing in homozygous *Abca4^ms1961G/G^*wild-type mice (Figure 3D). This allowed us to quantify base editing in a model, which presents two alleles that can be targeted by the mouse gRNA (ms-gRNA). At four weeks post-injection, A8 editing rates in the target cells were very similar to the rates obtained in the heterozygous *Abca4^hu1961E/ms1961G(KO)^*mice, with on average 61% and 85% editing on *Abca4* mRNA from retina and RPE/choroid/sclera, respectively (Supplemental Table 1, rows I, J, K and L). Thus, similarly to the heterozygous *Abca4^hu1961E/ms1961G(KO)^*model, our data suggest that the percentage of edited cells in models carrying two targetable alleles (*Abca4^ms1961G/G^* mice, wild-type NHPs, human retinal- and RPE/choroid explants) corresponds to the measured editing percentage.

Finally, we evaluated base-editing efficiency in NHPs, which are the most relevant animal models for macular diseases, since among mammalian models, only NHPs have a macula. We chose to deliver base editors via subretinal injection, which is a well-established delivery route for therapeutic agents to the retina^18, 19, 30^. We injected twelve adult Cynomolgus macaques with an equal mixture of AAV-SABE(N) and AAV-SABE(C) at three different dose levels (high dose = 5 × 10^11^ v.g./eye, mid dose = 3 × 10^11^ v.g./eye, and low dose = 1 × 10^11^ v.g./eye) (Figure 4A and Supplemental Table 4). We selected candidate base editor-vectors based on our in vitro results and we also considered other candidates with possible enhanced in vivo efficacy (Figure 4A). We injected 13 eyes with AAV5-SABE1 (high dose: 4 eyes, mid dose: 6 eyes, and low dose: 3 eyes). We injected four eyes with a high dose of a second candidate, AAV5-SABE2, which contains the W3-late SV40 polyA instead of bGH polyA. We chose this 3’UTR as the W3 element might confer an in vivo expression benefit in combination with the intronless CMV promoter in vivo^31^. We also injected four eyes with a chicken β-actin (CBA)-promoter-driven editor, AAV5-SABE3, at the high dose, as this promoter is part of the approved gene therapy product voretigene neparvovec^18^.

**Figure 4:**
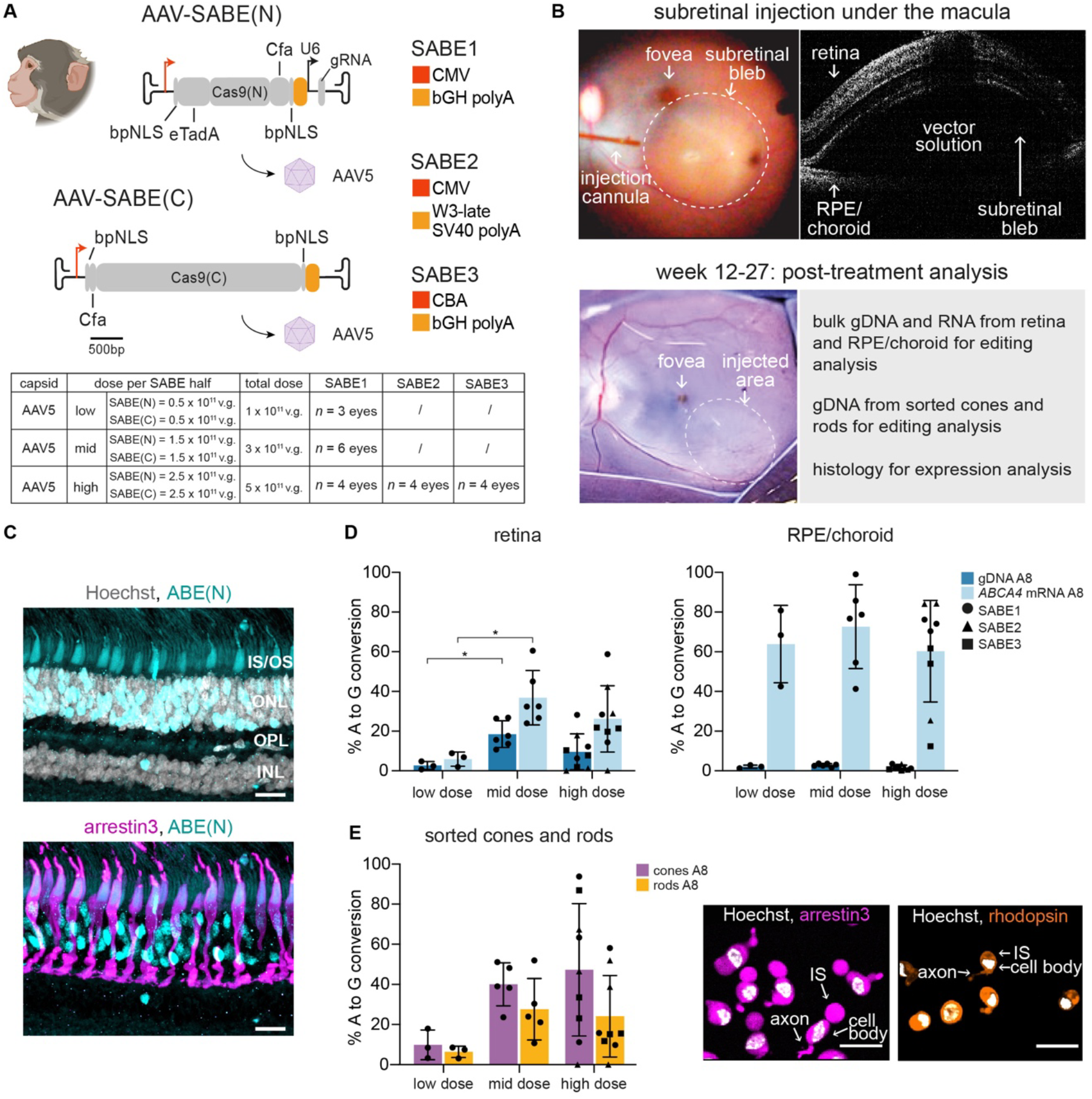
In vivo base editing in non-human primates. **A:**Experimental design. Schematic of the SABE(N) and SABE(C) constructs and the different dose levels used in the NHP study. **B:** Dual AAV5-SABE vectors were delivered to 21 eyes of NHPs by subretinal injection. OCT was used to confirm successful bleb formation. Eyes were harvested from 12- to 27 weeks post-injection and retina and RPE/choroid were processed separately for analysis. One half of the retinal tissue was used for bulk gDNA and RNA extraction, and the other half was dissociated for FAC-sorting of photoreceptors to determine editing rates in cones and rods. The edge of the injected area was used for histology. **C:** Representative immunofluorescence images of ABE(N) expression in a NHP retinal section (scale bars: 20 μm). IS/OS, photoreceptor inner and outer segments; ONL, outer nuclear layer (photoreceptors); INL, inner nuclear layer. Grey: Hoechst; cyan: ABE(N); magenta: arrestin3, gamma correction has been applied to obtain optimal dynamic range for visualization. **D:** In vivo base-editing efficiencies in gDNA and *ABCA4* mRNA with different base-editor constructs and at different doses. Results are presented as mean ± SD. Significance for dose response was calculated using a one-way ANOVA with Tukey’s correction, \**P* < 0.05. **E:** In vivo base-editing efficiencies in gDNA of sorted cones and rods and representative immunofluorescence images of sorted cells (scale bars: 25 μm). Results are presented as mean ± SD. White: Hoechst; magenta: arrestin3; orange: rhodopsin.

To confirm successful subretinal bleb formation, we used optical coherence tomography (OCT) immediately after surgery (Figure 4B). In three out of 21 eyes, OCT imaging showed no subretinal blebs and, therefore, these eyes were excluded from the study (Supplemental Table 4). Overall, the procedure was well tolerated and animals were kept for 12 to 27 weeks after injection. We then analyzed retinal- and the RPE/choroid tissues using histology and sequencing (Figure 4B). Immunostaining against ABE(N) revealed high expression of the base editor, mostly localized to the photoreceptor layer (Figure 4C). Editing at the gDNA and *ABCA4* mRNA in the retina was dose dependent, with the highest base editing observed at mid dose (an average of 18% editing in gDNA and 37% in *ABCA4* mRNA) (Figure 4D, left, and Supplemental Table 1, rows M and N). Editing rates at the RPE/choroid were consistently high at all the dose levels tested (on average 64% (low dose), 73% (mid dose), and 60% (high dose)) (Figure 4D, right and Supplemental Table 1, rows O and P). Editing rates in the retina and the RPE/choroid with the three different AAV5-SABE constructs were similar (Figure 4D, high dose and Supplemental Table 1, rows Q, R, S and T). To quantify editing rates in cones and rods, we sorted these cells by FACS and determined editing rates (Figure 4E and Supplemental Figure 12). Editing rates in cones were significantly higher than in rods (Supplemental Table 1, row U). At the mid dose we achieved on average 40% and 28% base editing rates in cones and rods, respectively. Altogether, the results show that the in vitro and mouse data translated to NHPs and that high levels of base correction were reached in vivo in the macaque eye after subretinal injection.

## DISCUSSION

In summary, we have demonstrated highly effective and precise base editing for Stargardt disease in the clinically relevant cell types of the human retina in vitro, and the mouse and NHP retina in vivo. In order to maximize the chance for successful clinical translation, we considered it important to establish and optimize our base-editing approach in human model systems first. Therefore, we validated our gene therapy in in vitro human model systems, such as retinal organoids and human-retinal and RPE/choroid explants, before testing candidate vectors in mice and NHPs in vivo. This approach resulted in effective translation of in vitro findings into in vivo base editing. A key requirement for therapeutic application of base editing is the demonstration of precise base correction in a high percentage of target cells. The evaluation of base-editing rates in *ABCA4* mRNA and from FAC-sorted photoreceptor cells, allowed us to specifically determine editing rates in cells expressing the causative gene. In the case of Stargardt disease, *ABCA4* is expressed in cone and rod photoreceptors and in RPE cells. At the optimal dose level, our optimized gene therapy vector led to on average 40% editing rates in cones, 28% in rods and 73% in RPE cells in NHPs in vivo at the A8 base, which we have demonstrated to be highly correlated to A7 target base editing. We found no off-target base editing in human retinal explants. These results suggest that adenine base editing is a rational therapy for correcting the most frequent mutation in Stargardt disease. The approaches we have developed will likely also be applicable to other neurodegenerative ocular diseases.

## SUPPLEMENTAL FIGURES AND SUPPLEMENTAL FIGURE LEGENDS

**Supplemental Figure 1:**
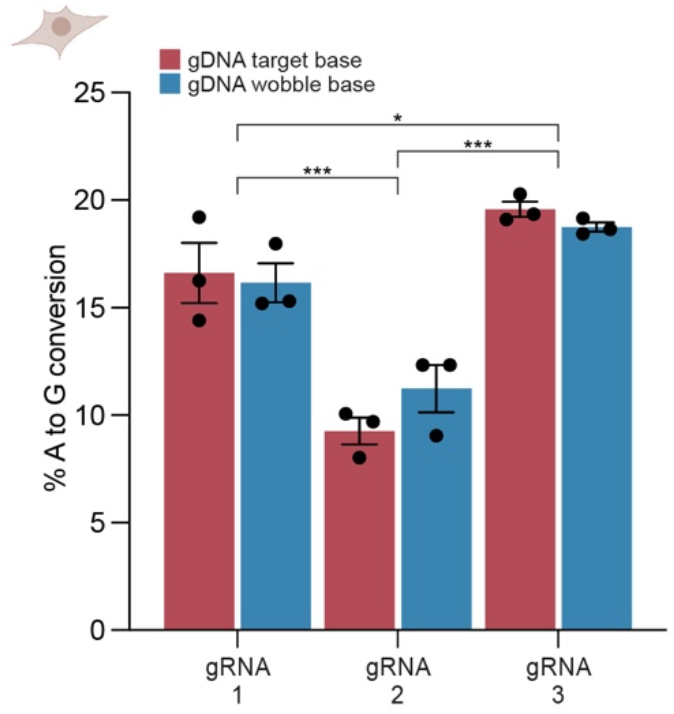
Adenine base-editing efficiency by different gRNAs. Base-editing efficiencies at the target- and wobble bases with the unsplit ABE7.10 base editor in combination with the different gRNAs. Results are presented as mean ± SD. *P < 0.05, ***P < 0.001 by three-way mixed-effect ANOVA with Tukey’s correction.

**Supplemental Figure 2:**
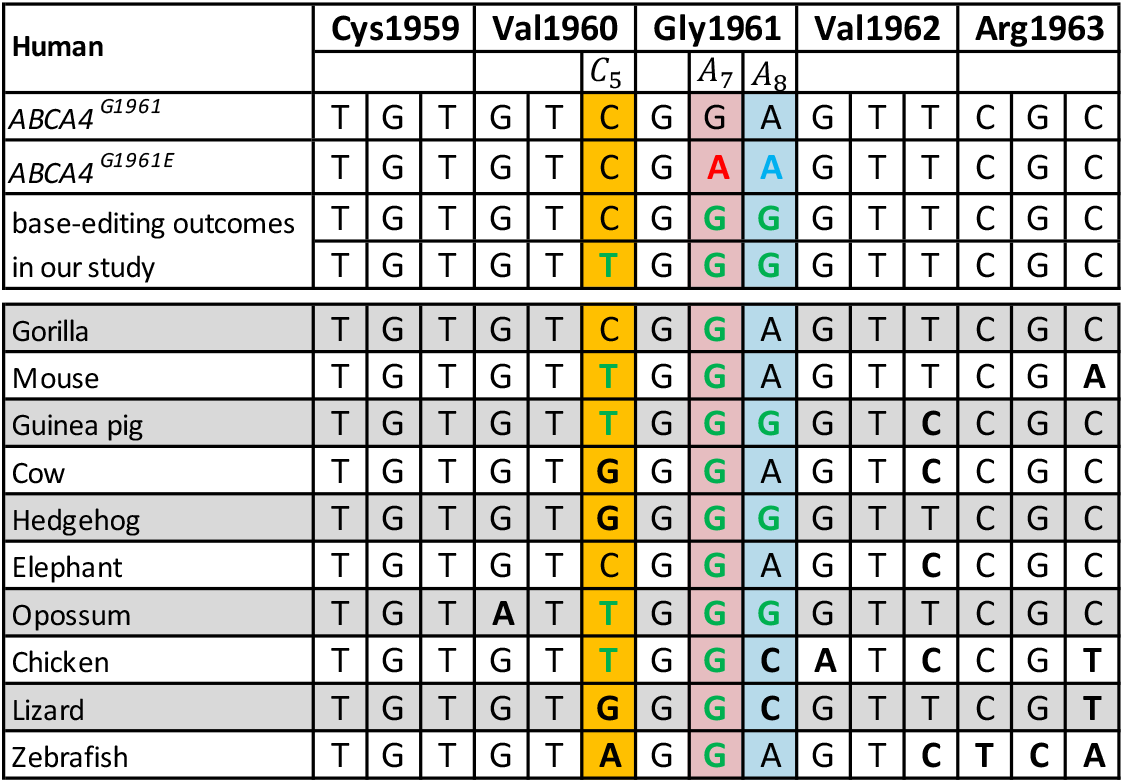
Conservation of the *ABCA4* sequence around p.Gly1961. The table shows the sequence alignment between humans and 10 other vertebrates. The first row shows the human *ABCA4* reference sequence. All sequence changes to the human sequence are indicated in bold. The second row shows the *ABCA4* c.5882A allele, with the A7 target base highlighted in red. The third and fourth rows show the most frequent base-editing outcomes in our study. The two observed bystander edits, c.5880C to c.5880T at position five (c.5880C>T, p.Val1960=) and c.5883A to c.5883G at position eight (c.5883A>G, p.Gly1961=) lead to silent changes, do not affect conserved base positions, and are present in other species. These results suggest that these bystander base changes have no biological relevance.

**Supplemental Figure 3:**
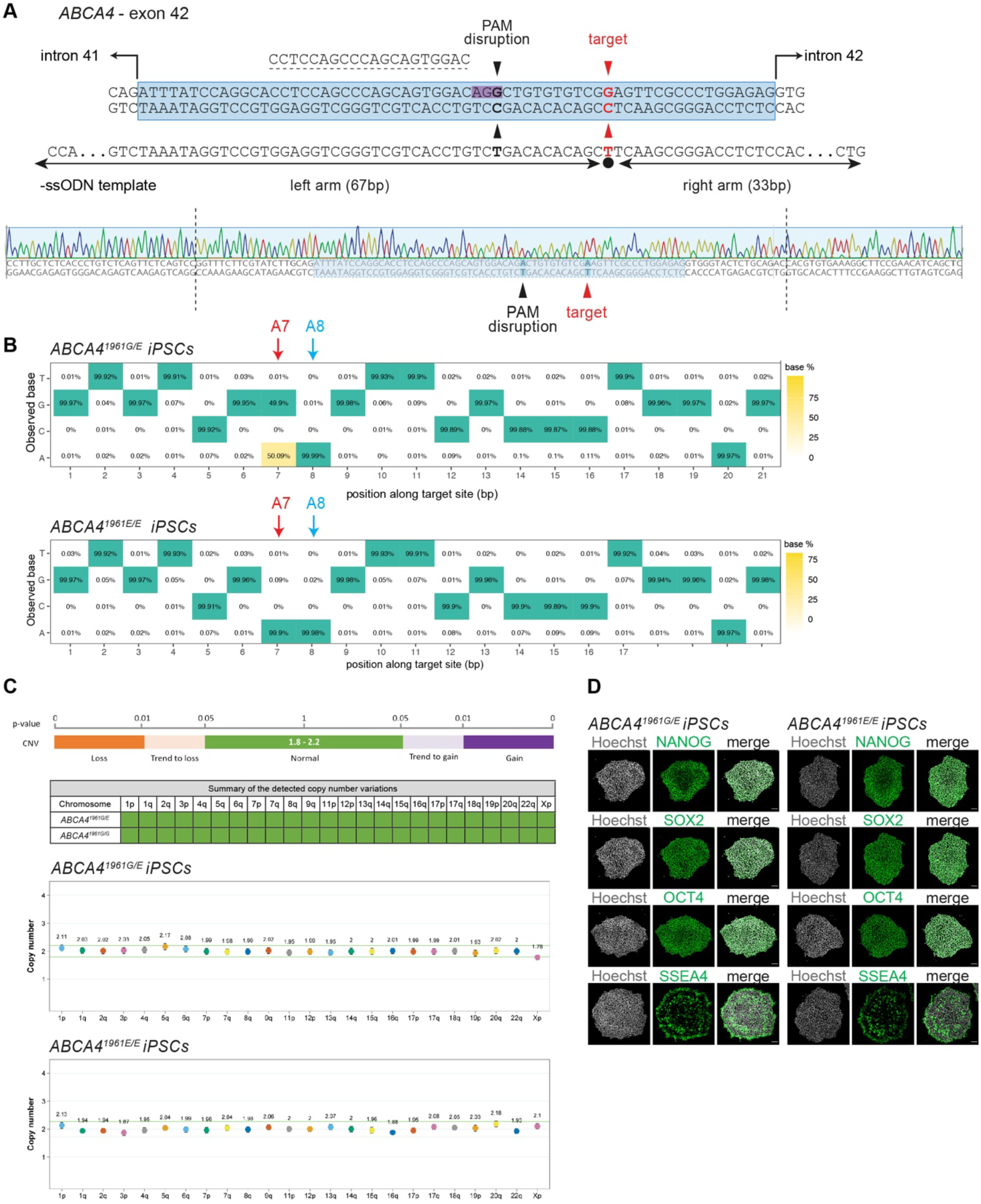
Generation of the *ABCA4^G1961E^* human mutant iPSC line. **A:**Strategy for the generation of an *ABCA4^G1961E^* iPSC line. *ABCA4* exon 42 (blue box) with flanking introns (top). The PAM site is highlighted in purple, and the gRNA binding site is indicated by a black dashed line. The black arrowheads point to the PAM disruption site (silent mutation) and the red arrowheads point to the *ABCA4* c.5882G>A mutation. Representative Sanger sequencing trace of the *ABCA4^1961E/E^* clone that was selected for human retinal organoid induction (bottom). **B:** Results from targeted deep-sequencing of the *ABCA4^1961G/E^* (top) and *ABCA4^1961E/E^* (bottom) clone confirming successful knock-in of the target mutation in a heterozygous or homozygous form. **C:** Results from the iPSC digital aneuploidy test, confirming the genomic integrity of the *ABCA4^1961G/E^* (top) and *ABCA4^1961E/E^* (bottom) clone. These clones were used for human retinal organoid induction. **D:** Confocal images of *ABCA4^1961G/E^* (left) and *ABCA4^1961E/E^* (right) iPSCs. Green: antibody for pluripotency markers (NANOG, SOX2, OCT4, SSEA4); grey: Hoechst (scale bars: 100 µm).

**Supplemental Figure 4:**
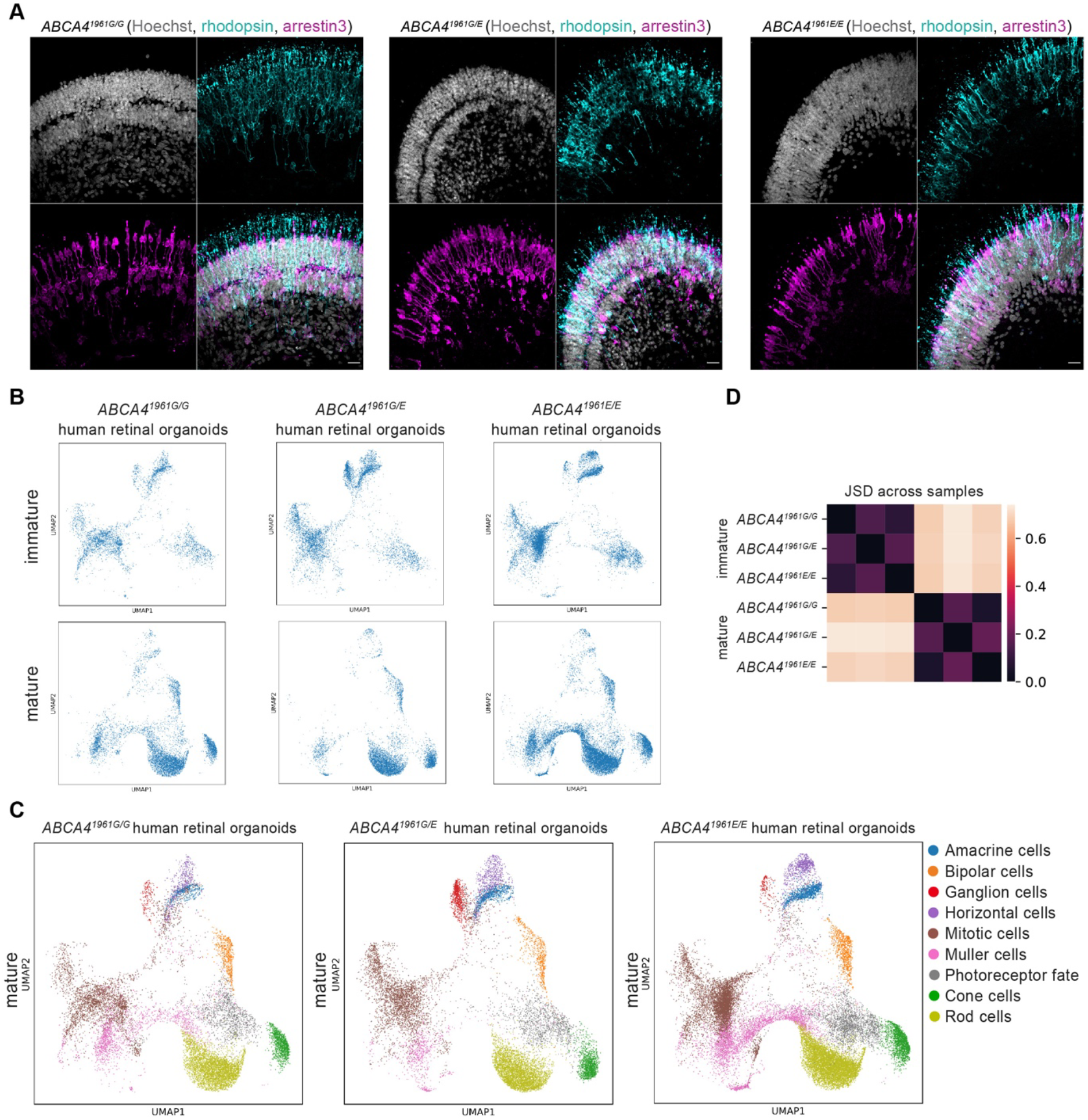
Characterization of the *ABCA4^G1961E^* human retinal organoids. **A:**Confocal images of *ABCA4^1961G/G^* (left), *ABCA4^1961G/E^* (middle), and *ABCA4^1961E/E^* (right). Grey: Hoechst; cyan: rhodopsin; magenta: arrestin3 (scale bars: 25 µm). **B:** 2D UMAP projection of single cells from human retinal organoids ordered by the *ABCA4* genotype (*ABCA4^1961G/G^* human retinal organoids: left; *ABCA4^1961G/E^* human retinal organoids: middle, and *ABCA4^1961E/E^* human retinal organoids: right) and the developmental stage (immature: top; mature: bottom). **C:** 2D UMAP plot of scRNA data from mature human retinal organoids colored by cell type and plotted separately by the *ABCA4* genotype (*ABCA4^1961G/G^* human retinal organoids: left; *ABCA4^1961G/E^*human retinal organoids: middle, and *ABCA4^1961E/E^* human retinal organoids: right). **D:** Heat map for Jensen–Shannon divergence (JSD) showing the similarity between organoids of different *ABCA4* genotypes at two different developmental stages (immature: top; mature: bottom).

**Supplemental Figure 5:**
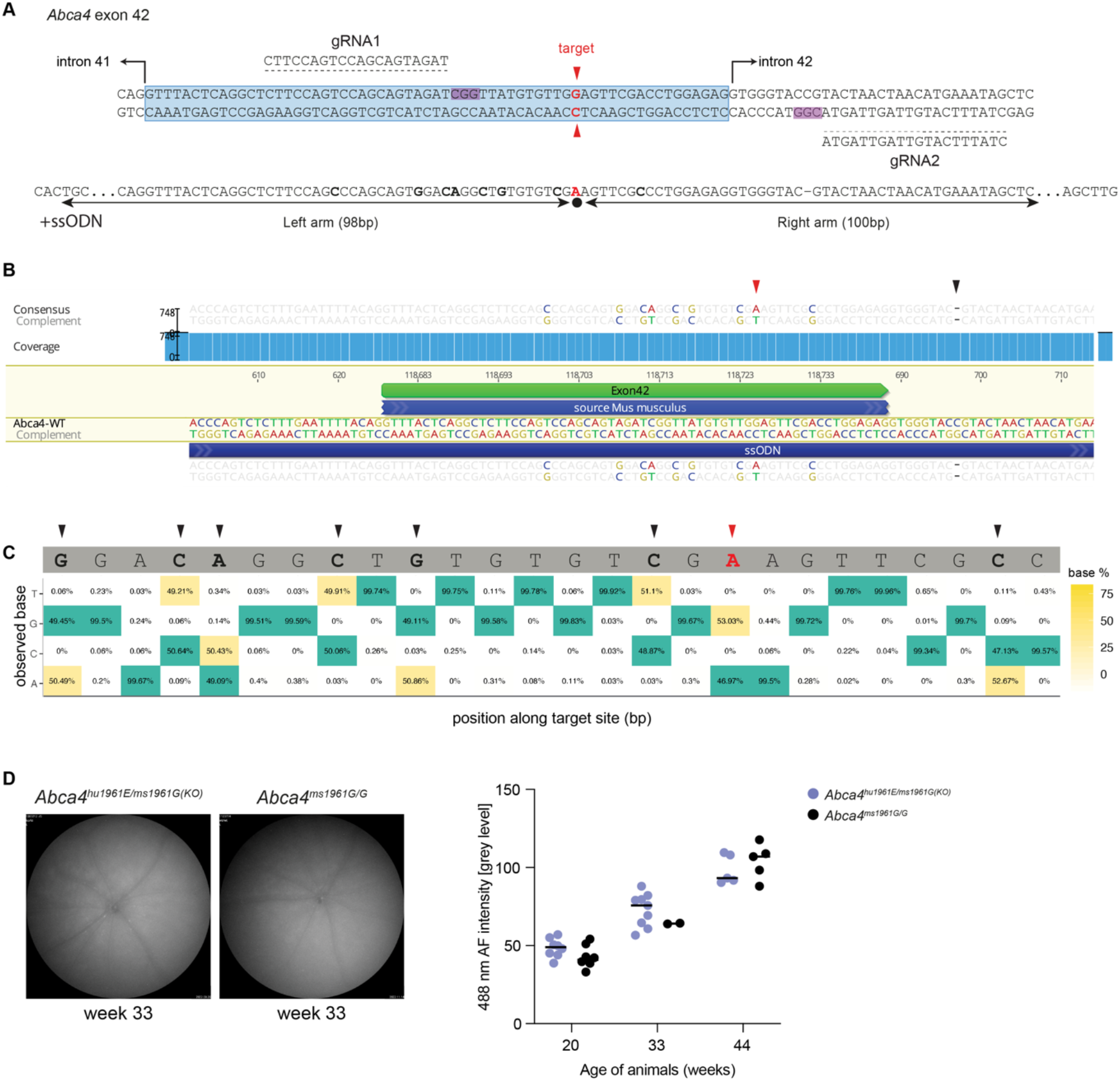
Generation of *Abca4^hu1961E^* mice. **A:**Strategy for the generation of the *Abca4^hu1961E^* mice line. *Abca4* exon 42 (blue box) with flanking introns. The PAM sites are highlighted, and the gRNA binding sites are indicated by black dashed lines. The red arrowhead points to the G1961E mutation. Bold nucleotides indicate nucleotide changes due to humanization. Note the deletion in the downstream intron – this is intentional and was introduced to disrupt the PAM site. The deletion is not expected to interfere with splicing as it is at position +9, at which there is no base preference for canonical splicing^32^. **B:** Sequencing of the gDNA of the *Abca4^hu1961E^* allele. The red arrowhead points to the *Abca4* c.5882G>A mutation, the black arrowhead points to a deletion in the intron. **C:** Deep-sequencing of *Abca4^hu1961E/ms1961G(KO)^* mice, where the results indicate heterozygosity. The red arrowhead points to the *Abca4* c.5882G>A mutation and the black arrowheads point to the nucleotide changes due to humanization. **D:** Fundus autofluorescence images of retinas from an *Abca4^hu1961E/ms1961G(KO)^* and wild-type *Abca4^ms1961G/G^* mouse (left). Quantification of the fluorescent signals at different ages (right).

**Supplemental Figure 6:**
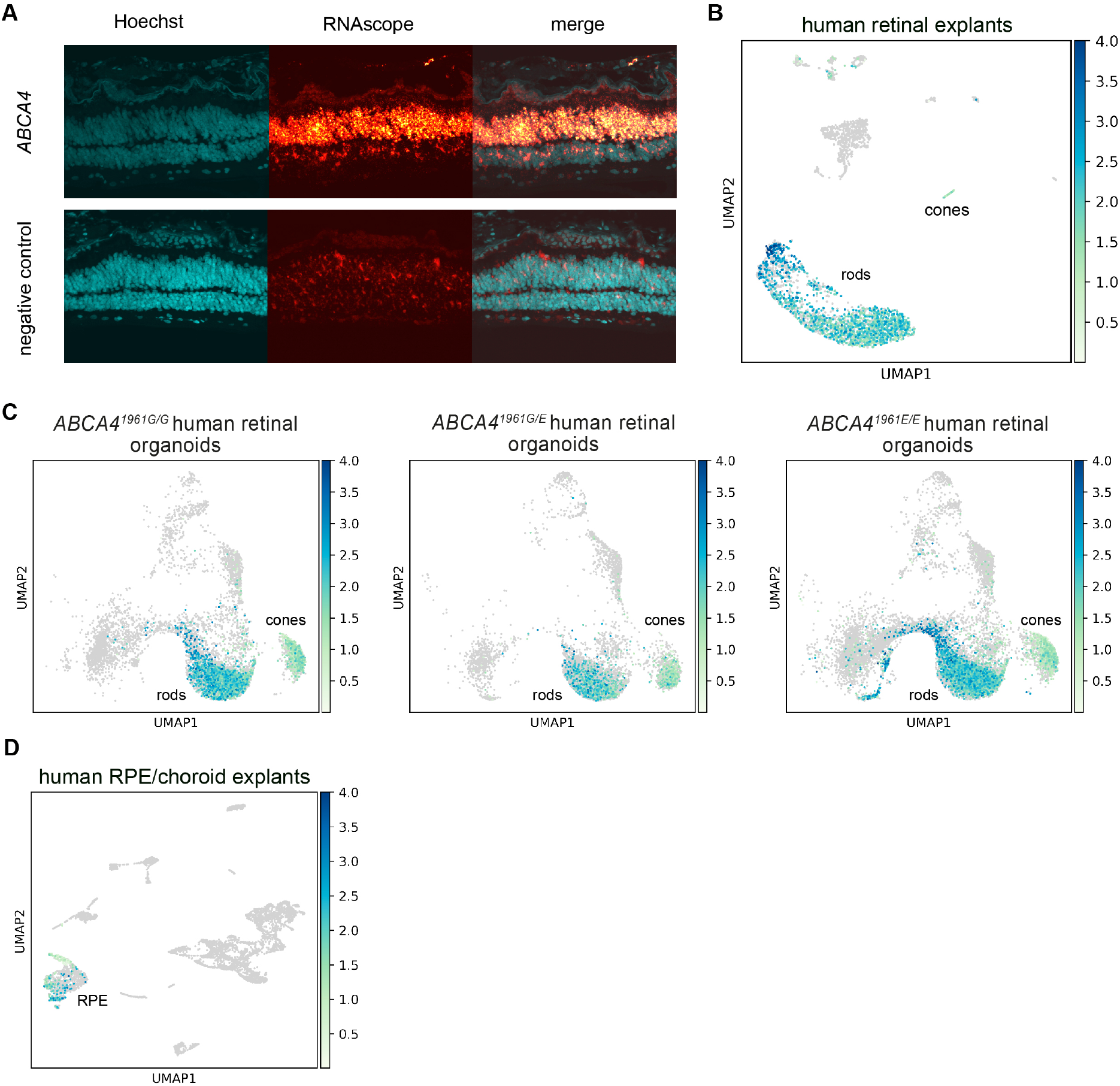
*ABCA4* expression in human model systems. **A:** RNAScope from human retina showing *ABCA4* expression in photoreceptors. **B-D:** UMAP clustering of scRNA data, highlighting *ABCA4-*expressing cells in blue. Results show that *ABCA4* is expressed in human cones, rods and RPE cells.

**Supplemental Figure 7:**
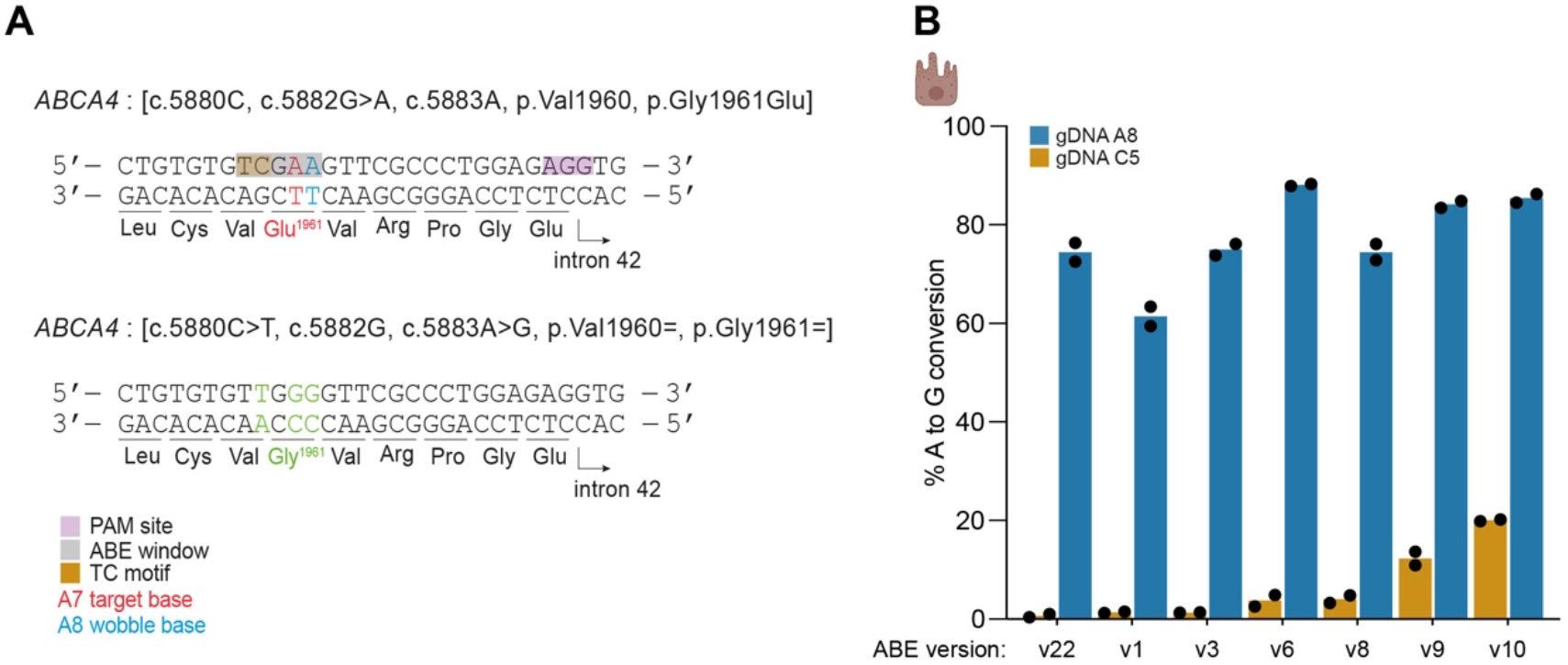
ABE-mediated cytosine editing on the *ABCA4* gene. **A:**Fragment of the *ABCA4* exon 42 with the base-editor window highlighted in grey. The base-editor window also contains a TC sequence that constitutes a possible motif for ABE-mediated cytosine editing. *ABCA4* c.5880C to c.5880T bystander editing results in a silent change (c.5880C>T, p.Val1960=). This change is not conserved and is expected to have no biological relevance (Supplemental Figure 2). **B:** Adenine base-editing efficiencies at the A8 site and cytosine base-editing efficiencies at the C5 site with different dual AAV-ABE versions in human iPSC-RPE.

**Supplemental Figure 8:**
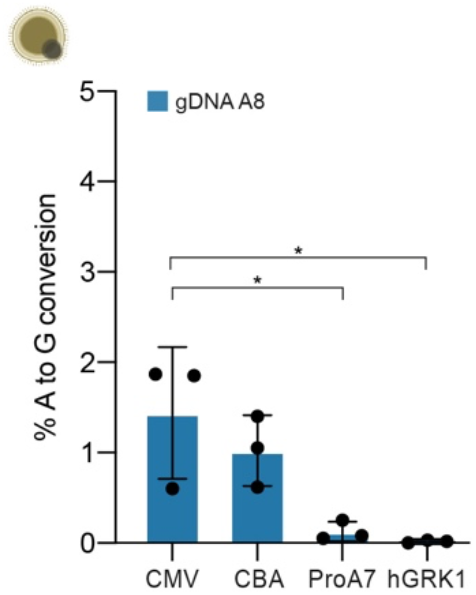
Testing of different promoters in *ABCA4^1961G/G^* human retinal organoids. Comparison of in vitro base-editing efficiencies at the A8 site in gDNA with different ubiquitous and photoreceptor specific promoters. Results are presented as mean ± SD. \**P* < 0.05 by one-way ANOVA with Tukey’s multiple comparisons test. CMV: cytomegalovirus promoter, CBA: chicken β-actin promoter, ProA7: cone-specific promoter from^17^, hGRK1: human rhodopsin-kinase promoter.

**Supplemental Figure 9:**
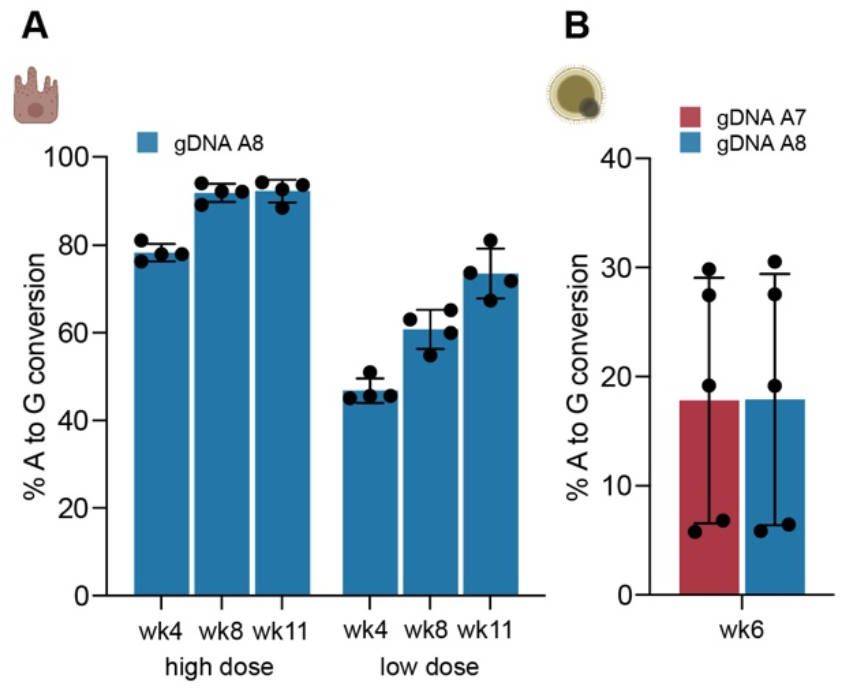
Dual AAV9-PHP.eB SABE1 base editing in different in vitro model systems. **A:**Base-editing efficiencies at the A8 site in gDNA of human iPSC-RPE at different time points and two different doses (high dose = 10^6^ v.g./cell and low dose = 10^5^ v.g./cell). **B:** Base-editing efficiencies at the A7 and A8 sites in gDNA of *ABCA4^1961E/E^* human retinal organoids. Results are presented as mean ± SD.

**Supplemental Figure 10:**
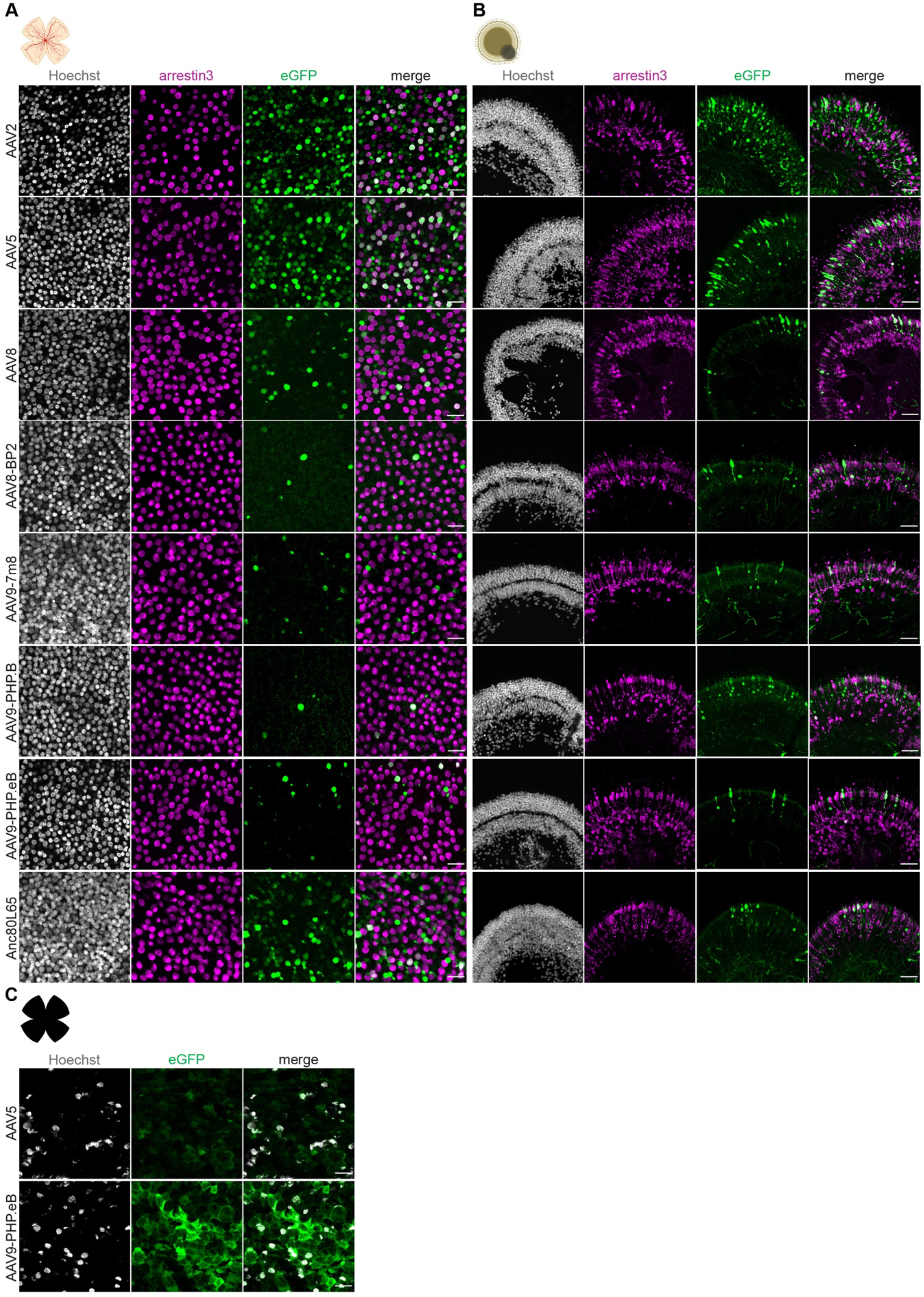
In vitro AAV capsid screen (low dose) **A:**Representative images of *ABCA4^1961G/G^* human retinal explants transduced with different AAV capsids encoding for CMV-eGFP. Results are from 5 weeks after transduction (4.7 × 10^10^ v.g./explant). Efficient cone-photoreceptor transduction is shown by colocalization of eGFP with arrestin3 (merge) (scale bars: 25 µm). **B:** Representative images of *ABCA4^1961G/G^* human retinal organoids transduced with the same capsids 4 weeks after transduction (3 × 10^10^ v.g./organoid). Efficient cone-photoreceptor transduction is shown by colocalization of eGFP with arrestin3 (merge) (scale bars: 50 µm). **C:** Representative images of *ABCA4^1961G/G^*human RPE/choroid explants transduced with AAV5- or AAV9-PHP.eB capsids 5 weeks after transduction (4.7 × 10^10^ v.g./explant) (scale bars: 25 µm). Grey: Hoechst; magenta: arrestin3; green: eGFP.

**Supplemental Figure 11:**
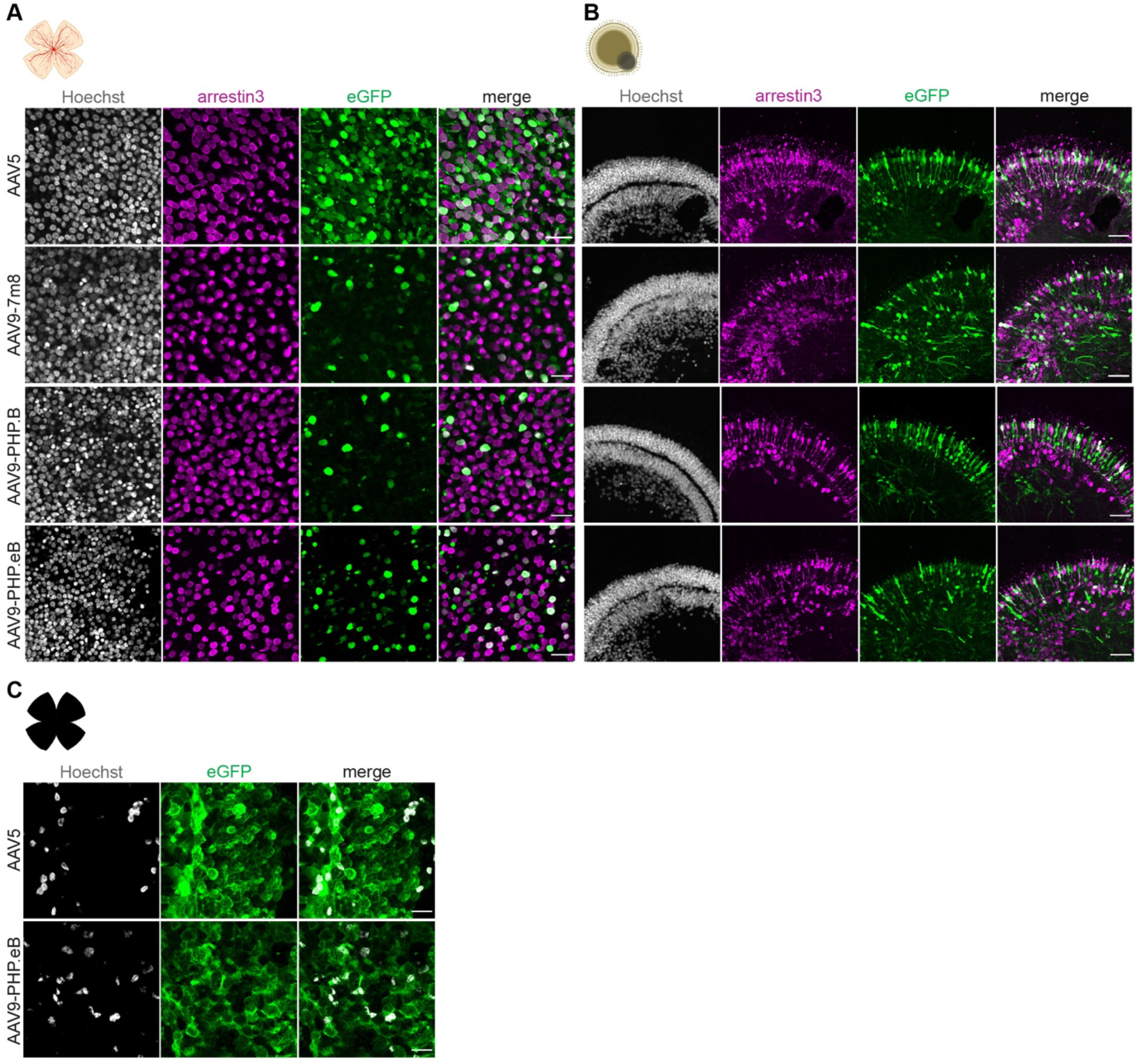
In vitro AAV capsid screen (high dose) **A:**Representative images of *ABCA4^1961G/G^* human retinal explants transduced with different AAV capsids encoding for CMV-eGFP. Results are from 5 weeks after transduction (2.5 × 10^11^ v.g./explant). Efficient cone-photoreceptor transduction is shown by colocalization of eGFP with arrestin3 (merge) (scale bars: 25 µm). **B:** Representative images of *ABCA4^1961G/G^* human retinal organoids transduced with the same capsids 4 weeks after transduction (1.15 × 10^11^ v.g./organoid). Efficient cone-photoreceptor transduction is shown by colocalization of eGFP with arrestin3 (merge) (scale bars: 50 µm). **C:** Representative images of *ABCA4^1961G/G^*human RPE/choroid explants transduced with AAV5- or AAV9-PHP.eB capsids 5 weeks after transduction (2.5 × 10^11^ v.g./explant) (scale bars: 25 µm). Grey: Hoechst; magenta: arrestin3; green: eGFP.

**Supplemental Figure 12:**
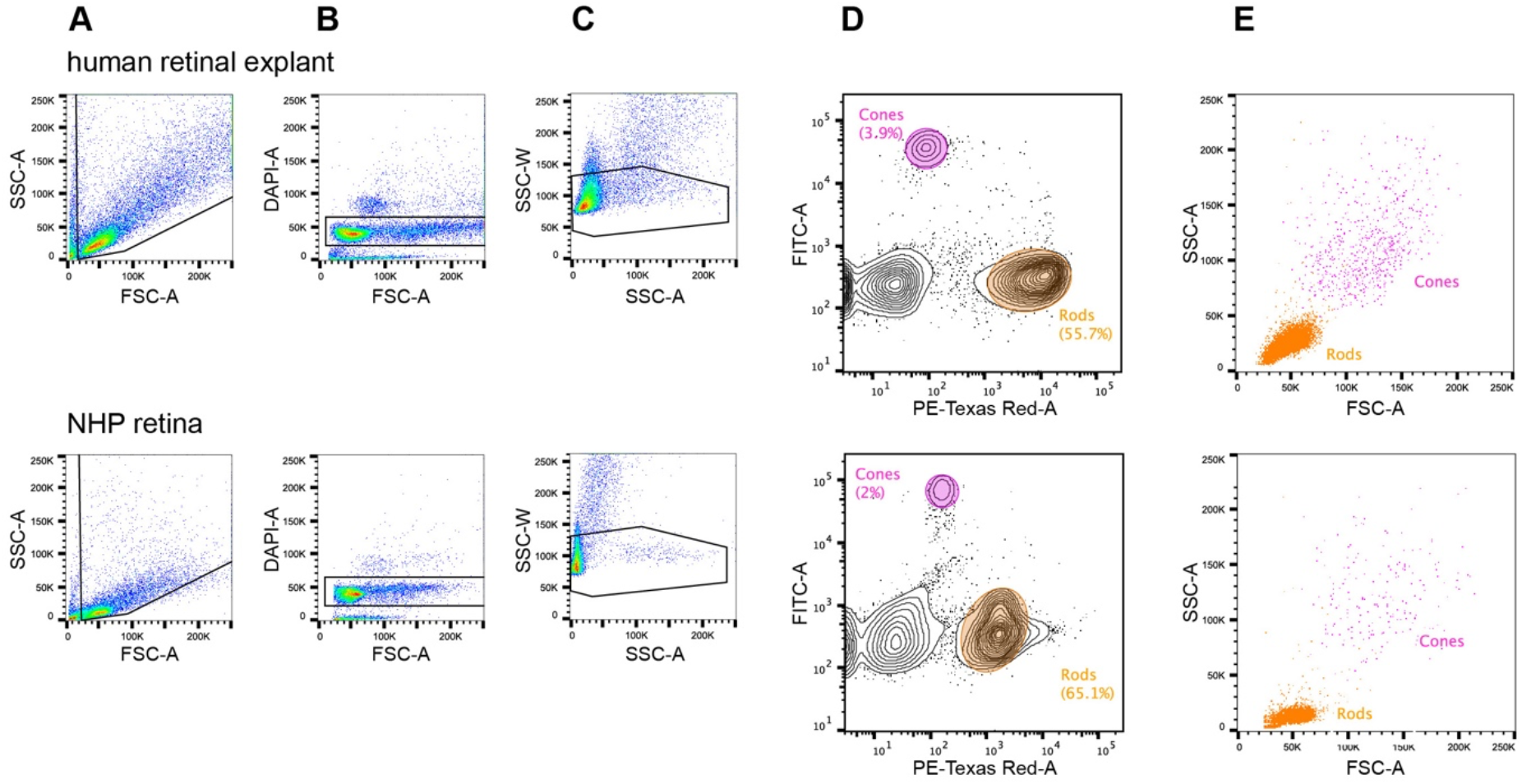
Isolation of cone- and rod-photoreceptor cells from human retinal explants and NHP retinas A-C: Representative FACS density plots from a human retinal explant (top) and a NHP retina (bottom) showing the initial gating strategy. **A:** The fixed cell suspension was first analyzed by forward scatter (FSC) and side scatter (SSC). **B:** To separate cells from debris, cells were further gated by Hoechst intensity. **C:** Cells were separated into singlets inside scatter area (SSC-A) versus height (SSC-H) plots. **D:** Contour plots depicting the two sorting gates based on arrestin3 (FITC-A+, cones) and rhodopsin (PE-Texas Red-A+, rods) expression and the frequencies of the two sorted populations. **E:** FACS dot plots showing sorted cones and rods by their size (FSC-A) and granularity (SSC-A), illustrating the expected size difference between the two photoreceptor cell types.

## SUPPLEMENTAL TABLES

**Supplemental Table 1:**
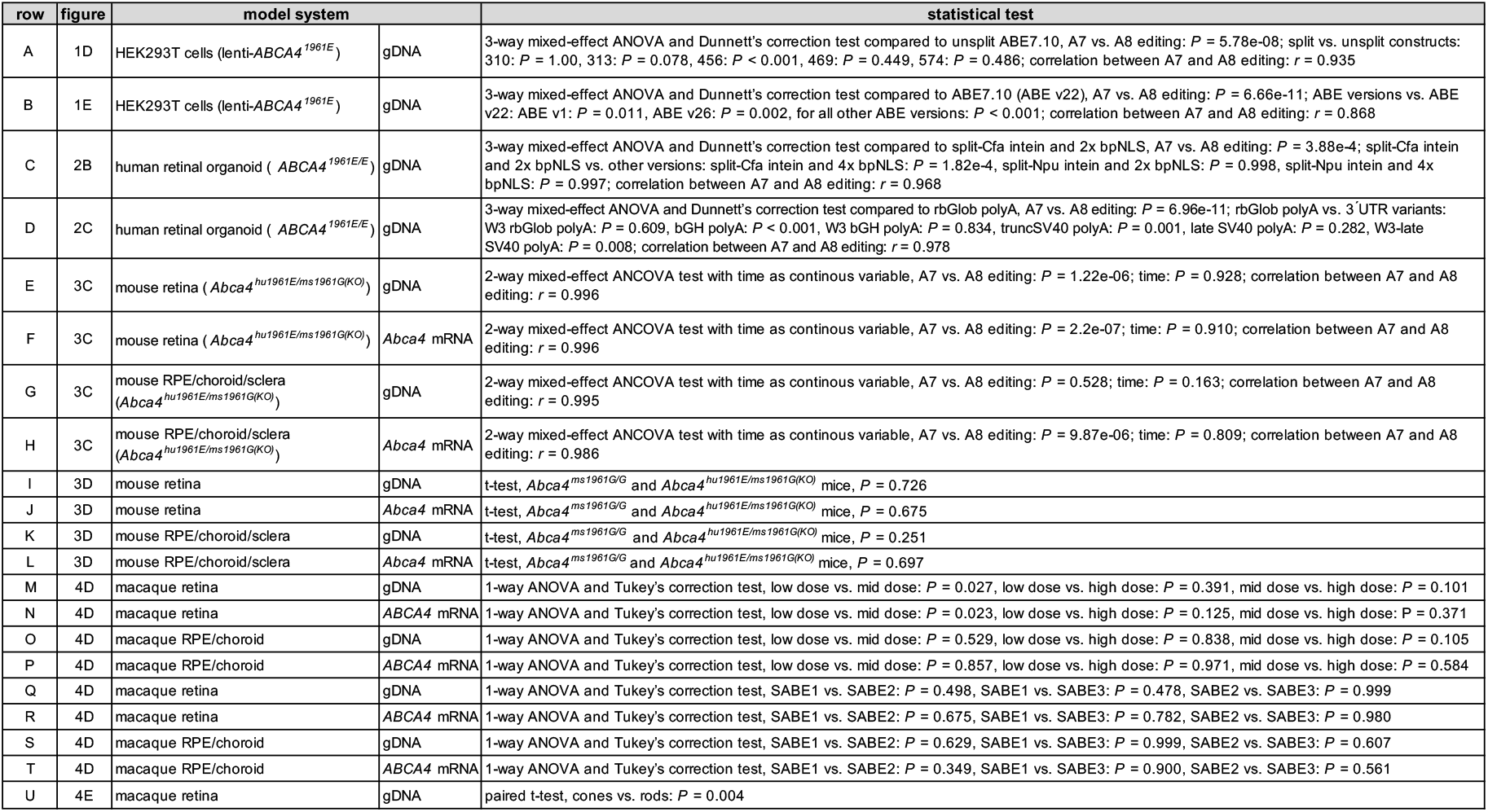
Details of statistical tests.

**Supplemental Table 2:**
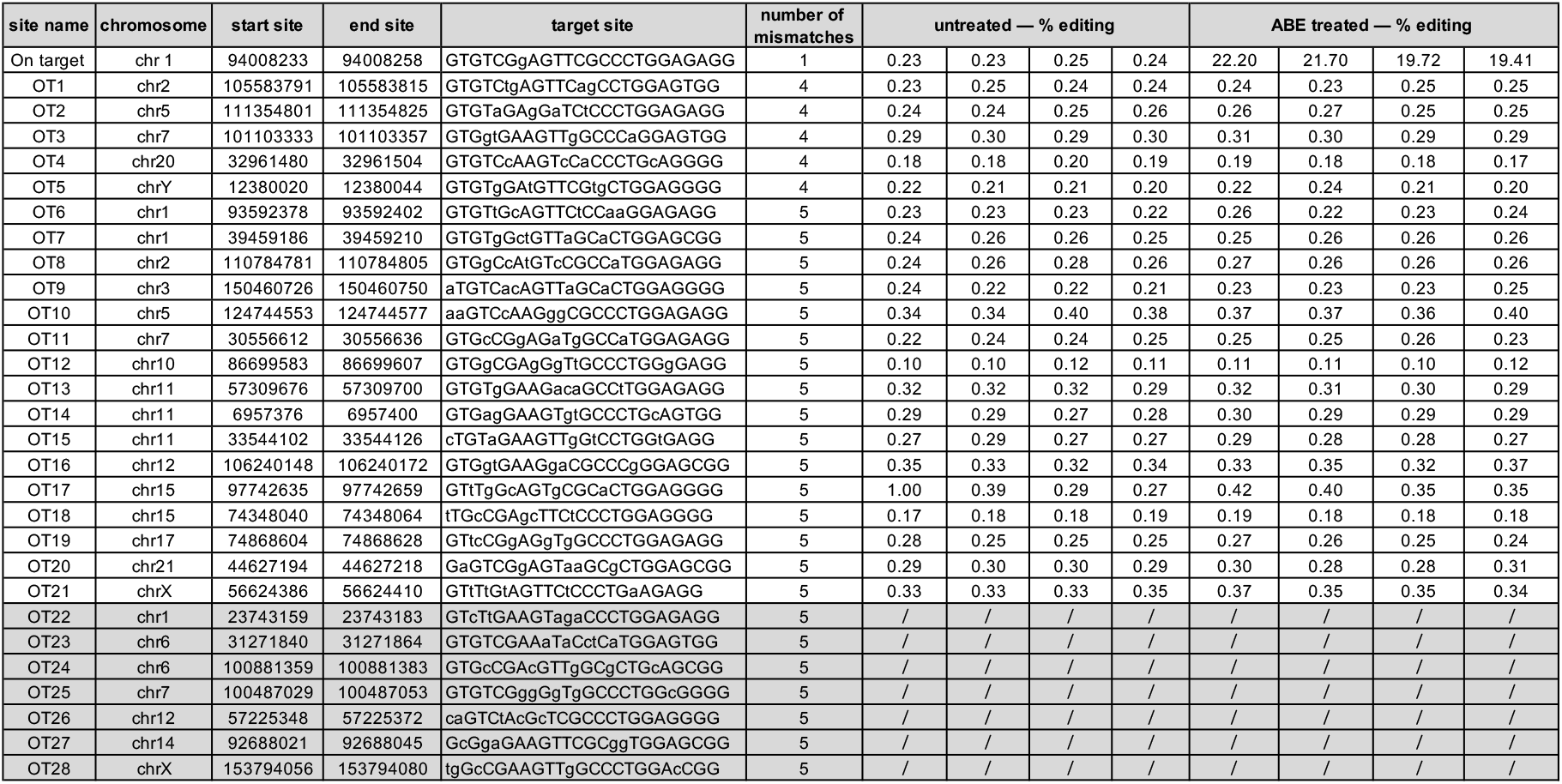
Details of off-target sites.

**Supplemental Table 3:**
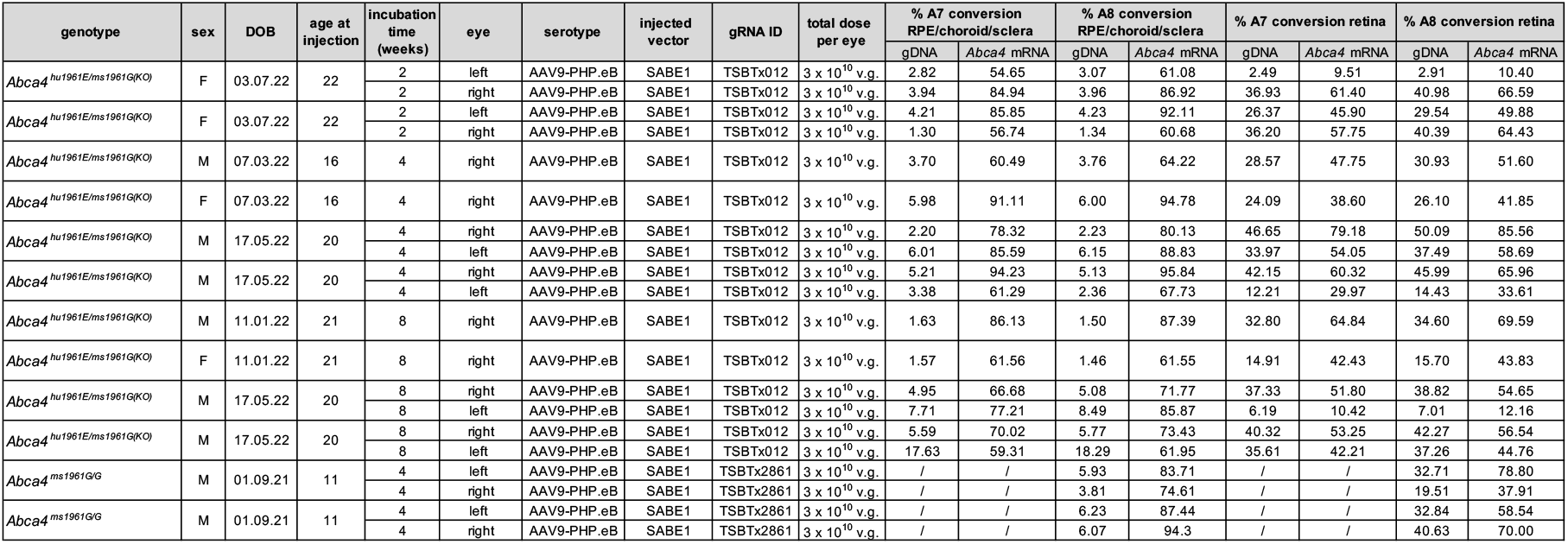
Detailed mouse study results.

**Supplemental Table 4:**
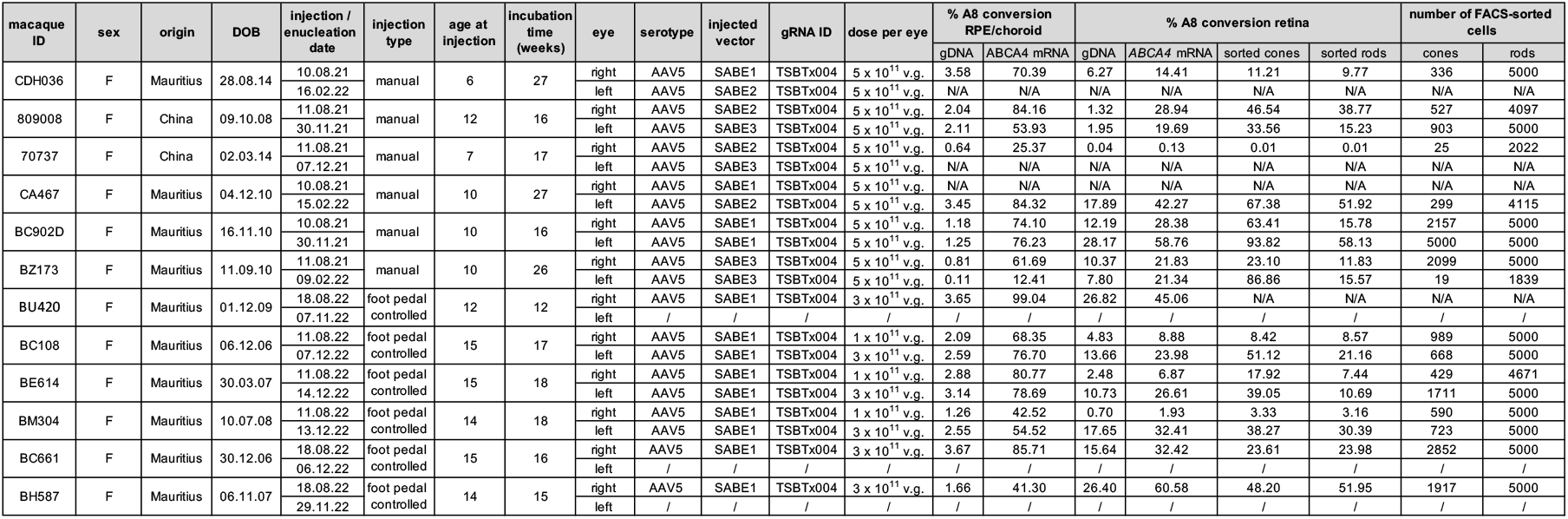
Detailed NHP study results.

**Supplemental Table 5:**
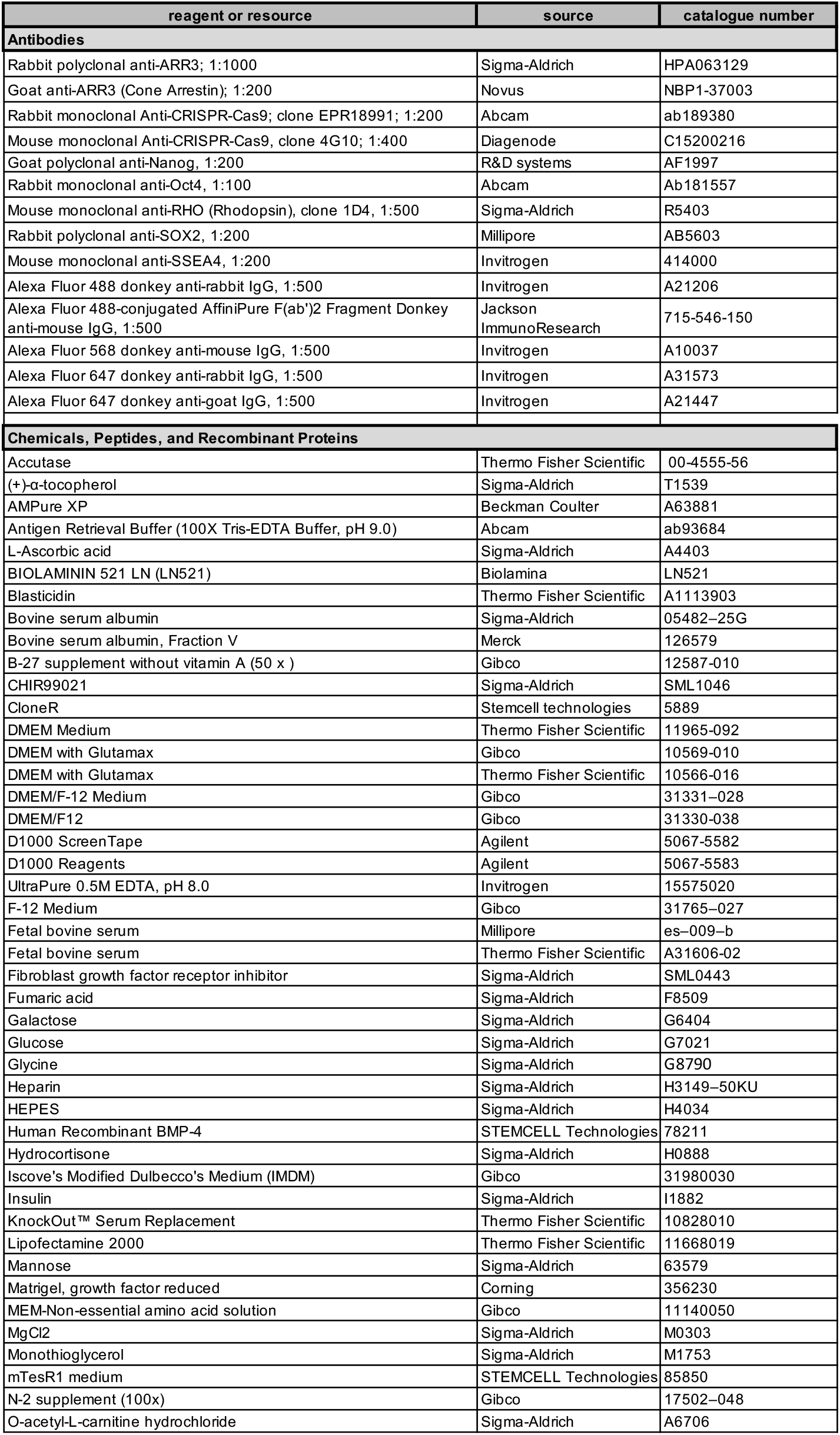
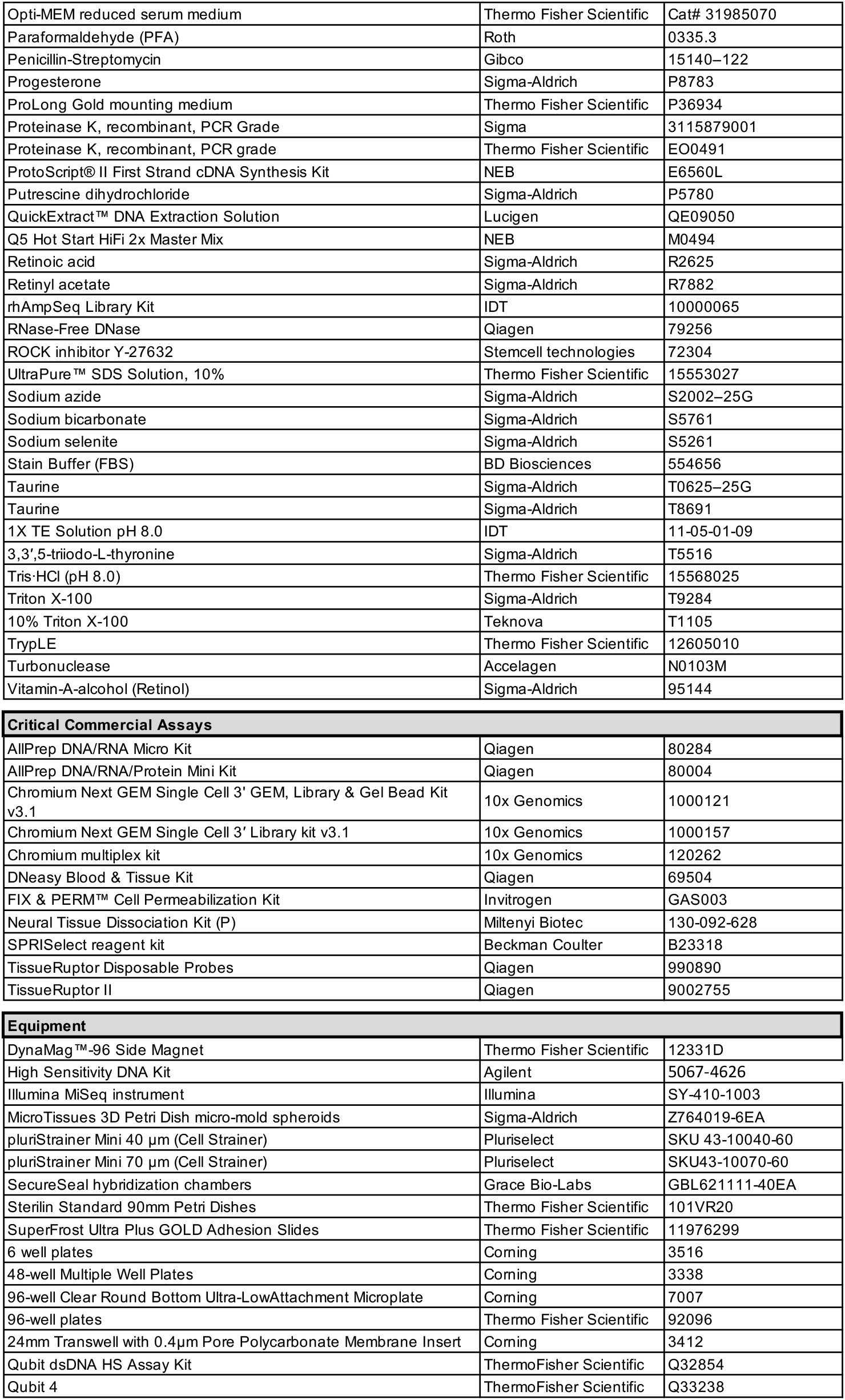
List of reagents, antibodies and biological samples.

## METHODS

Note: Reference numbers for reagents are found in Supplemental Table 5.

IIustrations of model systems in Figure 1C, D, and E, Figure 2B, C, D, E and H, Figure 3A, Figure 4A, Supplemental Figure 1, Supplemental Figure 7B, Supplemental Figure 8, Supplemental Figure 9A and B, Supplemental Figure 10 A, B and C and Supplemental Figure 11 A, B and C were created by BioRender (https://www.biorender.com/).

### Cell culture and generation of the HEK293T cell line containing a lentivirus-integrated *ABCA^41961E^*

HEK293T cells (American Type Cell Culture Collection, ATCC; CRL-3216) were cultured in Dulbecco’s Modified Eagles Medium (DMEM) plus Glutamax (Thermo Fisher Scientific) supplemented with 10% (v/v) fetal bovine serum (Thermo Fisher Scientific). A lentiviral production plasmid encoding the human V5-tagged *ABCA4* gene fragment, comprising the sequence from 72-bp upstream to 123-bp downstream of exon 42 and including the *ABCA4* c.5882G>A mutation, was generated by restriction cloning using a 5’ HpaI and 3’ ApaI restriction site flanked gBlock (IDT). The gene fragment was ligated into an HpaI/ApaI digested pLenti6.4 R4R2 V5-DEST vector (Thermo Fisher Scientific). Small-scale lentivirus was produced at Charles River Laboratories (Vigene Biosciences) and used to transduce HEK293T cells at a multiplicity of infection (MOI) of 0.3–10 infectious units (IFU)/cell. Cells harboring stable lentivirus integrants were selected by 10 µg/mL blasticidin (Thermo Fisher Scientific) and maintained in the presence of 5 µg/mL blasticidin. The average number of lentiviral integrations per cell was assessed by targeted amplicon sequencing using primers oBTx361 and oBTx362, which simultaneously amplify the virally integrated *ABCA4* fragment and the endogenous *ABCA4* locus. The number of viral integrations in each sample was estimated by multiplying the number of NGS reads containing the *ABCA4* c.5882G>A mutation by two and dividing by the number of wild-type reads (integration number = mutant reads × 2 / wt reads). Cell lines containing an average of two or fewer integrations per cell were used for all base-editing experiments.

### Base editing in HEK293T cells

Base-editor plasmids were codon optimized and synthesized by GeneArt (Thermo Fisher Scientific). The base editor was intein-split to two halves in the SpCas9 sequence at amino acid positions 310, 313, 456, 469 and 574 (the number refers to the first amino acid on the ABE(C)). The first amino acid on ABE(C) was mutated to cysteine. Lenti-*ABCA4^1961E^*HEK293T cells were seeded in Corning CellBIND 48-well Multiple Well Plates (Corning) at a density of 35,000 cells per well in DMEM supplemented with Glutamax and 10% (v/v) FBS, but without blasticidin. Cells were transfected ∼24 h after seeding. Complementary plasmid pairs containing the split-intein ABE and guide RNA (gRNA), or a full-length base-editor plasmid and gRNA, were combined at a 1:1 molar ratio (1,000 ng) for transfection with 1.5 μL Lipofectamine 2000 (Thermo Fisher Scientific) in a total volume of 25 µL Opti-MEM reduced serum medium (Thermo Fisher Scientific). The reagent mixtures were added to the wells following the manufacturer’s instructions. Media was replaced every 48 h over a 5-day period before cell lysis. After 48 h, media were removed and cells washed twice with 100 µL of phosphate buffered saline (PBS, Thermo Fisher Scientific) before adding 75 µL of cell lysis buffer (10 mM Tris·HCl; pH 8.0) (Thermo Fisher Scientific) + 0.05% SDS (Thermo Fisher Scientific) + 100 µg/mL Proteinase K (Thermo Fisher Scientific). Wells were scraped with pipette tips and the cell lysate immediately transferred to a 96-well plate. The plate was incubated at 55°C for 1 h followed by heat inactivation at 95°C for 20 min. Samples were then stored at −20°C before further processing.

### AAV vector production and titration

Recombinant AAV vectors were produced using transient triple transfection of suspension cultures of HEK293T cells with plasmids containing (i) AAV Rep and Cap genes, (ii) the transgene flanked by inverted terminal repeat (ITR) sequences, and (iii) adenovirus genes (E4, E2a and VA) necessary for AAV replication (helper plasmid). Cells were lysed 72 h after transfection and unpackaged DNA removed by adding Triton X-100 (Teknova), MgCl_2_ (Sigma-Aldrich), and Turbonuclease (Accelagen) at final concentrations of 0.25% (v/v), 2 mM, and 10 U/mL, respectively. Cell lysis was performed at 37°C for 2-5 h in a shaker incubator. The cell lysate was then filtered through a clarification depth filter, followed by filtration through a 0.2-µm filter. Clarified lysate was loaded onto the affinity chromatography column of a ÄKTA Pure chromatography system (Cytiva). Captured AAV was eluted using an elution buffer at pH 2.5-3. The eluate was captured and the pH promptly neutralized. Full and empty AAV particles were separated by cesium chloride density gradient ultracentrifugation. Bands containing full AAV particles were collected and the titer determined by qPCR. The full particle samples were then diluted in CsCl stock solution to a final target concentration and dialyzed into AAV formulation buffer (10 mM Na_2_HPO_4_, 2 mM KH_2_PO_4_, 2.7 mM KCl, 192 mM NaCl, 0.001% Pluronic F-68; pH 7.4) using 100-kDA cutoff dialysis cassettes (Thermo Fisher Scientific). Dialysate was filtered using low-protein binding 0.1-µm syringe filters and then aliquoted and stored at –80°C. Final AAV titers were determined using droplet digital PCR (ddPCR) from vialed material. Other quality attributes such as endotoxin level, aggregation status, osmolality, and pH were also determined.

### Generation of an iPSC line containing the *ABCA4* c.5882G>A mutation using CRISPR- Cas9

We used CRISPR-mediated homology-directed repair (HDR) to knock-in the *ABCA4* c.5882G>A mutation into the previously published 01F49i-N-B7 iPSC line^33^ (Supplemental Figure 3). An additional silent change (c.5871G>A) was introduced to inactivate the protospacer adjacent motif (PAM) site. A two-part gRNA complex was formed by combining the designed crRNA (Alt-R CRISPR-Cas9 crRNA, IDT) with the tracrRNA (Alt-R CRISPR- Cas9 tracrRNA, IDT). The gRNA was combined with the Cas9 nuclease protein (Alt-R SpCas9 Nuclease, IDT) to form a CRISPR-Cas9 ribonucleoprotein complex (RNP complex). The repair template containing the target mutation and the PAM disruption was designed as a 100- bp-long single-stranded oligonucleotide (ssODN) with asymmetric homology arms and complementarity to the non-target strand (Supplemental Figure 3). ssODN was ordered at a 100-µM scale from Sigma-Aldrich. At 60%–70% confluency, 01F49i-N-B7 iPSCs were dissociated to single cells and 250,000 cells transfected with 25 μL of the prepared transfection mix containing 20 µL of nucleofection buffer (P3 Primary Cell 4D-NucleofectorTM X Kit S, Lonza), 5 µL of the RNP complex, 1.2 µL of ssODN template, and 2.6 µL PBS. The transfection was performed in a 4D-Nucleofector X Unit (Lonza) using the CA-137 program. After transfection, 75 μL of 37°C pre-warmed mTesR1 medium (Stemcell technologies) supplemented 1:10 with CloneR (Stemcell technologies) was added to the cuvette and the cells transferred to a 6-well plate. Cells were kept in culture until 80-90% confluency. For identification of successfully edited clones, individual colonies were picked manually and expanded in a 96-well plate. Genomic DNA (gDNA) was extracted by adding 40 µL QuickExtract DNA Extraction Solution (Lucigen) to each well. After incubation at room temperature for 10 min, wells were scraped with pipette tips and lysates transferred to PCR tubes. The tubes were incubated at 68°C for 15 min followed by 95°C for 8 min. We performed targeted PCR by adding 1 μL of cell lysate (1:5 dilution) to a 25-μL PCR reaction containing GoTaq Hot Start Master Mix Green (Promega) and 0.5 μL of the primers (10 µM, oT01-24 and oT01-25). PCR reactions were carried out as follows: 95°C for 3 min, 34 cycles of (95°C for 30 s, 55°C for 30 s, and 72°C for 2 min), and a final 72°C extension for 10 min. To characterize edited iPSC clones, we performed long-range Sanger sequencing, targeted deep-sequencing, flow cytometry to analyze pluripotency markers, and an aneuploidy test using ddPCR (Stemgenomics) (Supplemental Figure 3).

### Single-cell RNA sequencing

Wild-type *ABCA4^1961G/G^*, heterozygous *ABCA4^1961G/E^*and homozygous *ABCA4^1961E/E^* human retinal organoids from the 01F49i-N-B7 iPSC line were subjected to single-cell sequencing. We also single-cell-sequenced human retinal- and RPE/choroid explants. Cells were dissociated as described below. We loaded 8,000 cells onto a 10x Genomics Chromium Next GEM Chip G and ran it in the Chromium Controller to generate single-cell Gel Beads in Emulsion (GEMs). Single-cell RNA-Seq libraries were prepared using the Chromium Next GEM Single Cell 3’ Reagent Kits (10x Genomics) version 3.1 according to the manufacturer’s manual (version CG000204_Rev_C for the version 3.1 kit). Reverse transcription of GEMs was performed at 53°C for 45 min followed by 85°C for 5 min. After reverse transcription, the GEMs emulsion was broken and the first-stranded complementary DNA (cDNA) cleaned-up with DynaBeads MyOne Silane Beads. cDNA was amplified at 98°C for 3 min, 12 cycles of (63°C for 20s and 72°C for 1 min), with a final incubation at 72°C for 1 min. The amplified cDNA product was then cleaned-up with the SPRIselect Reagent Kit (0.6 × SPRI; Beckman Coulter). Indexed sequencing libraries were constructed using the reagents in the Chromium Next GEM Single Cell 3′ Library kit v3.1 (10x Genomics) and following these steps: (i) fragmentation, end repair and A-tailing, (ii) double-sided size selection with SPRIselect Reagent Kit (0.6 × SPRI and 0.8 × SPRI), (iii) adaptor ligation, (iv) clean-up with SPRIselect (0.8 × SPRI), (v) sample indexing PCR by the Chromium multiplex kit (10x Genomics), and (vi) double-sided size selection with SPRIselect Reagent Kit (0.6 × SPRI and 0.8 × SPRI). The barcode sequencing libraries were quantified using the Qubit 2.0 with a Qubit dsDNA HS assay kit (Thermo Fisher Scientific) and the quality of the libraries assessed on a 2100 Bioanalyzer (Agilent, Santa Clara, CA, USA) using the High Sensitivity DNA Kit (Agilent, Santa Clara, CA, USA). Sequencing libraries were loaded at 10 – 12 pM on an Illumina HiSeq2500 with 2 × 50 paired-end kits using the following read length for the Chromium v3.1 kit: 28 cycles for Read1, 8 cycles i7 index, and 91 cycles for Read2.

### iPSC-derived retinal pigment epithelial (iPSC-RPE) cell culture and AAV transduction

Human iPSC-RPE cells were differentiated with small modifications according to previous reports^34^. Briefly, embryoid bodies (EBs) were generated from the iPS(IMR90)-4-DL-01 line^33^ by seeding 9,000 single cells per well in agarose microwell arrays (Sigma-Aldrich). EBs were cultivated in 6-well plates (Corning) in growth factor-free chemically defined medium (gfCDM) containing Iscove’s Modified Dulbecco’s Medium (IMDM, Thermo Fisher Scientific), 45% F12 (Thermo Fisher Scientific), 450 µM monothioglycerol (Sigma-Aldrich), 1% penicillin/streptomycin (Thermo Fisher Scientific), and 10% KnockOut Serum Replacement (Thermo Fisher Scientific). Six days post EB induction, 1.5 nM bone morphogenetic protein 4 (BMP4, Stemcell technologies) was added to the medium for another three days. Half of the medium was then exchanged with fresh gfCDM every three days. At day 18, RPE spheroids were transferred to a 10-cm cell culture dish and kept in suspension in ‘1:1 medium’ containing DMEM: F12 (Thermo Fisher Scientific), 1% N2 Supplement (Thermo Fisher Scientific), 1% penicillin/streptomycin, 5 µM fibroblast growth factor receptor inhibitor (FGFR, Sigma-Aldrich) and 3 µM GSK inhibitor (CHIR99021, Sigma-Aldrich). The medium was exchanged every three days until the RPE spheroids were fully pigmented (12-15 days). Thereafter, to obtain a homogeneous monolayer of 2D RPE, RPE spheroids were dissociated into single cells with TrypLE (Thermo Fisher Scientific) and seeded on Matrigel-coated (Corning) 96-well plates (Corning) at a density of 100,000 cells/cm2. Cells were cultured in 1:1 medium supplemented with 10% FBS (Millipore) until confluency, and maintained in 1:1 medium with 1% FBS. RPE cells were transduced with AAV vectors in a 96-well plate at a multiplicity of infection of 1 × 10^5^ or 1 × 10^6^ vector genomes (v.g.) per cell for each split-intein ABE half and maintained in 50 µL of 1:1 medium at 37°C in 5% CO2. After 4 h, 50 µL of media was added to each well. One day later, 100 µL of media was added to each well. After 24 h, and every 48 h thereafter, the solution was completely exchanged with fresh media. Four to 11 weeks later, samples were either fixed or we proceeded with gDNA isolation as described below.

### Human retina and RPE/choroid culture and AAV transductions

Human retinas and RPE/choroid samples were collected at the Department of Ophthalmology, Semmelweis University (Budapest, Hungary) or at the Department of Ophthalmology, Basel University Hospital (Basel, Switzerland). All tissue samples were obtained in accordance with the tenets of the Declaration of Helsinki. Personal identifiers were removed and samples were coded before processing. All experimental protocols were approved by the local ethics committees (ethical permit numbers: Budapest: ETT TUKEB 34851-2/2018/EKU and ETT TUKEB IV/5645-1/2021/EKU, Basel: EKNZ 2021-01773). Briefly, after enucleation, the vitreous was removed and the retina and the RPE/choroid layers were separated from each other. We cultured ∼5 × 5 mm tissue pieces on polycarbonate membrane inserts (Corning) with the photoreceptor side down or the choroid side down. The cultures were maintained at 37°C in 5% CO_2_ in DMEM/F12 medium (Thermo Fisher Scientific), supplemented with 0.1% BSA (Merck), and (from Sigma-Aldrich) 10 μM O-acetyl-L-carnitine hydrochloride, 1 mM fumaric acid, 0.5 mM galactose, 1 mM glucose, 0.5 mM glycine, 10 mM HEPES, 0.05 mM mannose, 13 mM sodium bicarbonate, 3 mM taurine, 0.1 mM putrescine dihydrochloride, 0.35 μM retinol, 0.3 μM retinyl acetate, 0.2 μM (+)-α-tocopherol, 0.5 mM ascorbic acid, 0.05 μM sodium selenite, 0.02 μM hydrocortisone, 0.02 μM progesterone, 1 μM insulin, 0.003 μM 3,3′,5-triiodo-L-thyronine, 2,000 U penicillin, and 2 mg streptomycin. The media was changed every second day. AAV vectors containing eGFP or base editors were added 1-2 days after enucleation at 1.66 × 10^11^ v.g./explant. Four to seven weeks later, samples were either fixed or we proceeded with bulk- gDNA and RNA isolation as described below. For flow cytometry, cells were dissociated as described below.

### Mice

Mice were used in accordance with standard ethical guidelines as stated in the European Communities Guidelines on the Care and Use of Laboratory Animals. All animal experiments and procedures were approved by the local ethics committee (permit number: 3048/31896, Kantonales Veterinäramt Basel-Stadt). All mice were maintained in a pathogen-free environment with ad libitum access to food and drinking water. C57BL/6J wild-type mice were obtained from Charles River Laboratories. *Abca4^huG1961E/E^* mice (B6-*Abca4^em1(huEx42-G1961E)Brsk^*, short: *Abca4^hu1961E/E^*) were created by the University of Basel, Center for Transgenic Models (Basel, Switzerland). Briefly, we used the CRISPR-EZ technique^35^ with two gRNAs (Supplemental Figure 5) to humanize 8 nucleotides in a 34 nucleotide-long region in exon 42 of the *Abca4* gene and to simultaneously introduce the *Abca4* c.5882G>A mutation. One-cell embryos were electroporated with SpCas9/gRNA RNP particles and an ssODN template. One C nucleotide in the +9 position in intron 42 was deleted to inactivate the PAM site. This nucleotide deletion is not expected to alter splicing and is not conserved between mouse and human. The mouse line was validated using primers that bind outside of the targeted region (oT06-37 and oT06-38). Targeted deep-sequencing was used to confirm the sequence identity (Supplemental Figure 5). To obtain *Abca4^hu1961E/ms1961G(KO)^* heterozygous mice, we crossed *Abca4^huG1961E/E^* animals to B6.129S-Abca4^tm1Ght^/J mice (JAX:026800, Jackson Laboratories, Bar Harbor, ME, USA).

### Subretinal injection of mice

Subretinal injections were performed at age 11-22 weeks (Supplemental Table 3) as described before^17^. In brief, animals were anesthetized by subcutaneous injection of fentanyl, medetomidine and midazolam (0.05 mg/kg, 0.5 mg/kg, 5 mg/kg, respectively). A small incision was made with a sharp 27 G needle at the corneal-scleral divide and a total of 3 × 10^10^ v.g./eye of AAV mixture or 1.5 μL of formulation buffer was injected through this incision into the subretinal space using a blunt 5 μL Hamilton syringe (Hamilton Company) held by a micromanipulator. Animals were recovered by subcutaneous injection of naloxone, atipamezole and flumazenil (1.2 mg/kg, 2.5 mg/kg, 0.5 mg/kg), respectively.

### Processing of mouse eyes

After sacrifice, eyes were removed using curved forceps (Fine Science Tools) and transferred into ice cold 1X HBSS (Thermo Fisher Scientific) for dissection. Extraocular muscle and optic nerve were carefully removed using spring scissors (Fine Science Tools), followed by dissection of the cornea and lens. The retina was then carefully separated from the RPE/choroid/sclera and processed separately for bulk gDNA and RNA isolation or fixed for histology (see below).

### NHPs

Twelve healthy female Cynomolgus macaques (*Macaca fascicularis*, aged 6- to 15 years old; 10 animals from Mauritius and 2 from China) were used in the study (Supplemental Table 4). Animals were housed at the Simian Laboratory Europe facility (Silabe, Niederhausbergen, France), monitored in accordance with the guidelines of the European Directive 2010/63, and handled in strict accordance with good animal practice as defined by the French National Charter on the Ethics of Animal Experimentation. All animal protocols were approved by the French Ministry of Higher Education and Research (permit number: APAFIS#27357- 2020092811266511_v2 (28/12/2020)).

### Subretinal injections in macaques

Animals underwent a comprehensive ophthalmological examination by a veterinary ophthalmologist before injection. We obtained pre-injection fundus photos and OCT scans of the macula and the optic nerve head confirming that the eyes were healthy. Also, NHPs were genotyped to exclude polymorphisms in the target region via PCR / Sanger sequencing using oT04-46, oT04-47, oT04-48, oT04-49. Three days before injection, treatment of the animals with 0.75 mg/kg intramuscular dexamethasone began and continued for one week. One day before injection, animals received 15 mg/kg intramuscular amoxicillin followed by two further doses 48 h apart. Animals were fasted before the day of the surgery (but access to water was maintained). On the day of the surgery, animals were anaesthetized using 10 mg/kg intramuscular ketamine and transported to the operating room. The pupils were dilated using eyedrops containing 0.5% tropicamide and 10% phenylephrine. In the operating room, animals received propofol (5-10 mg/kg) followed by intubation. The anesthesia was maintained by isoflurane (1-2.5%). The first 6 animals were injected through a two-port vitrectomy configuration and a manual injection. After disinfection of the ocular surface, two ports were made on the limbus using 25G trocars, one for the endoillumination port and one for the 41G subretinal microinjection cannula. After gently touching the retina with the subretinal cannula, a slow manual injection was performed with the BSS to induce a pre-bleb. The second 6 animals were injected through a three-port vitrectomy configuration and the injection was controlled via a foot-pedal. In this second cohort, a core vitrectomy was performed, followed by pre-bleb formation by BSS. Immediately after visible confirmation of the pre-bleb, the injection was stopped, and the virus solution was injected into this pre-bleb through the same retinotomy in the 12 animals. We injected 5 × 10^11^ v.g./eye, 3 × 10^11^ v.g./eye or 1 × 10^11^ v.g./eye from an equal mixture of the base-editor halves in a 70-µL target bleb volume. Shortly after injection, we performed OCT imaging to visualize the subretinal blebs. In three out of 21 injections, we found no OCT evidence for bleb formation, hence these eyes were excluded from the study (Supplemental Table 4). Animals received subconjunctival antibiotic immediately after the procedure and tobramycin ointment for 7 days following the procedure.

### Processing of NHP eyes

NHP eyes were removed during terminal anesthesia. Immediately after removal, the anterior segment including the cornea was removed by circularly cutting behind the limbus. The vitreous fluid was carefully removed by forceps and the eyecup processed further. Briefly, the bleb region was cut in half by an incision with a scalpel from the periphery towards the fovea. The central bleb was then punched out using a 4-mm biopsy punch. The retina and the underlying RPE/choroid were separately processed. One half of the retinal tissue of the punched region was used for dissociation and cell sorting (see below), while the remaining retina and the entire RPE/choroid were subjected to bulk- gDNA and RNA isolation. The circular edge of the tissue was fixed with 4% PFA in PBS and processed for histology. A control, non-treated area was also removed and processed similarly.

### Genomic DNA and RNA extraction from human iPSC-RPE cells and tissues

Genomic DNA from human iPSC-RPE was isolated by adding 40 µL QuickExtract DNA Extraction Solution (Lucigen) to each well. After incubation at room temperature for 10 min, wells were scraped with pipette tips and lysates transferred to PCR tubes. The tubes were incubated at 68°C for 15 min followed by 95°C for 8 min. Bulk gDNA and RNA from human retinal organoids were isolated using the AllPrep DNA/RNA Micro Kit (Qiagen) or DNeasy Blood & Tissue Kit (Qiagen) according to the manufacturer’s instructions. Bulk gDNA and RNA from human, mouse and macaque retina or RPE/choroid tissue were isolated using the AllPrep DNA/RNA Mini Kit (Qiagen). Briefly, tissue samples were simultaneously disrupted and homogenized using TissueRuptor II (Qiagen) for 20-30 s at full speed. During RNA purification, DNase digestion was performed on the column membrane using DNase I (Qiagen) for 15 min at room temperature according to the manufacturer’s instructions. RNA was eluted in 14-30 µL of RNase-free water. The elution step was repeated with the eluate. For gDNA, a one-step elution was performed in 50-100 µL of elution buffer (preheated to 70°C). Complementary DNA was synthesized from 20-60 ng of RNA using the ProtoScriptII First Strand cDNA Synthesis Kit (NEB).

### Fluorescence-activated cell sorting (FACS) of rods and cones

Retinal tissues were dissociated using the Neural Tissue Dissociation Kit (P) (Miltenyi Biotec). Briefly, 990 µL of enzyme mix P (40 µL enzyme P and 950 µL buffer X) was added to a ∼ 5 × 5 mm piece of human retina or a ∼ 4 × 3 mm piece of NHP retina and tissues incubated for 20-30 min at 37°C while shaking. Next, a further 10 µL of enzyme P was added and tissues incubated for another 10 min at 37°C. When dissociation was complete, 15 µL of enzyme mix A (5 µL enzyme A and 10 µL buffer Y) was added to the samples and mixtures were incubated for another 10-15 min at 37°C. The cell suspension was then centrifuged at 300 × *g* for 5 min at 4 °C and cells resuspended in 100 µL of stain buffer (BD Biosciences, supplemented with 5 mM EDTA (Thermo Fisher Scientific)). Cells were strained through a 70-µm filter (pluriSelect) followed by a 40-µm filter (pluriSelect) and fixed using 100 µL of fixation reagent (Medium A, Thermo Fisher). After 15 min, the cells were washed and resuspended in 300 µL of stain buffer. To stain cones and rods, samples were incubated in 100 µL of Medium B (Thermo Fisher Scientific) containing primary antibodies (arrestin3, Sigma-Aldrich, HPA063129, 1:33 and rhodopsin, Sigma-Aldrich, R5403, 1:100) for 30 min. Cells were then washed in stain buffer and secondary antibody staining (Alexa Fluor 488 donkey anti-rabbit IgG, A21206, Invitrogen, 1:100 and Alexa Fluor 568 donkey anti-mouse IgG A10037, Invitrogen, 1:100) was performed in 50 µL of stain buffer. To differentiate nucleated cells from debris, Hoechst was added at 1:165 dilution together with the secondary antibodies. After a 30- min incubation at room temperature in the dark, cells were washed twice with stain buffer and resuspended in 200-300 µL of stain buffer. A FACSAria (BD Biosciences) sorter was used to collect rods and cones into 20 µL of QuickExtract DNA Extraction Solution (Lucigen). The numbers of sorted cells ranged from 200- to 750 cones and 2,000- to 6,500 rods for human retinas (Figure 2G). For macaque retinas, the numbers of collected cells were 20- to 5,000 cones and 1,800- to 5,000 rods (Figure 4E and Supplemental Table 4). Immediately after sorting, proteinase K was added for reverse crosslinking (Sigma-Aldrich) and gDNA extraction performed (incubation at 56°C for 30 min, 65°C for 15 min, and 95°C for 10 min).

### Target amplicon sequencing DNA

For deep-sequencing of the *ABCA4* locus, we performed targeted PCR from gDNA, cDNA and cell lysates from human iPSC-RPE or sorted cells. Briefly, 2- to 10-µL gDNA samples or 5- µL cDNA samples were added to a 50-μL PCR reaction. Cell lysates (2 µL human iPSC-RPE or 10 µL sorted cells) were added to a 100-µL PCR reaction. The PCR reaction contained Q5 Hot Start HiFi 2X Master Mix and 0.4 μM of each primer containing 5’ Illumina adapter overhangs. PCR reactions were carried out at 95°C for 2 min, 30 cycles of (95°C for 15 s, 65°C for 20 s, and 72°C for 20 s), and a final 72°C extension for 2 min. Following amplification, 2 µL of the crude PCR products containing the amplified site of interest were barcoded using 0.5 µM of each unique Illumina barcoding primer pair and Q5 Hot Start High-Fidelity 2X Master Mix in a total volume of 25 µL. The reactions were carried out as follows: 98°C for 2 min, 10 cycles of (98°C for 20 s, 60°C for 30 s, and 72°C for 30 s), and a final 72°C extension for 2 min. Equal volumes of barcoded PCR products were then pooled and cleaned-up using SPRISelect paramagnetic beads (Beckman Coulter) using a 0.6 × bead/sample ratio. The amplicon was sequenced with an Illumina MiSeq instrument according to the manufacturer’s protocol.

### Targeted deep-sequencing analysis

FASTQ files were first generated from base call files (BCF) created by the MiSeq instrument using Illumina blc2fastq (v2.20.0.422). Adapters were trimmed using trimmomatic (v0.39) with parameters set up to clip Illumina TruSeq adapters, exclude reads shorter than 20 bases, trim the remaining 3’ end of reads if the average base quality (Phred score) in a 4-bp sliding window dropped below 15, trim any bases with quality scores of 3 or lower at the end of reads, and trim the 4 randomized bases introduced from the round 1 PCR primers. Trimmed reads were aligned to the reference amplicon sequence using bowtie2 (v2.35) in end-to-end mode with the --very-sensitive flag specified. The SAM files created by bowtie2 were converted to BAM files, sorted, and indexed using samtools (v1.9). BAM files were processed using the bam-readcounts tool (https://github.com/genome/bam-readcount) to generate plain text files summarizing the number of non-reference bases (substitutions), deletions, and insertions at each position in the alignment. The minimum base quality (Phred score) for counting a non-reference base was set to 29 in order to exclude low confidence base calls. Editing rates for each position in the target site were calculated as the fraction of non-reference bases of a given type (e.g., G) to the total number of bases passing the base-quality threshold at a given position in the alignment.

### Off-target screen

Initial in silico identification of candidate off-target sites was performed by running Cas- OFFinder v2.4^26^ on the GRCh38 human reference genome to identify genomic sequences harboring an NGG- PAM and up to 5 mismatches to the STGD-gRNA. For rhAmpSeq panel design, the list was further filtered; off-target sites with 5 mismatches to the STGD-gRNA were only included if they overlapped with an annotated gene exon (as annotated in GENCODE v24). The final custom rhAmpSeq CRISPR Panel (IDT) included a pool of 28 off-target sites and the c.5883A (A8) on-target site (Supplemental Table 2). PCR1 was set up with 5 µL 4X rhAmpSeq Library Mix 1 (IDT), 2 µL 10X rhAmpSeq forward panel, 2 µL 10X rhAmpSeq reverse panel, and 11 µL gDNA at 9.1 ng/µL. The PCR reaction was carried out as follows: 95°C for 10min, 10 cycles of (95°C for 15 s, 62°C for 4 min), and finally 99.5°C for 15 min. PCR products were cleaned-up using 1.5 × Ampure XP beads (Beckman Coulter, Brea, CA, USA): 30 µL of room temperature Ampure beads were added to each reaction, mixed, and incubated for 10 min at room temperature. Solutions were then exposed to a magnet (Thermo Fisher Scientific) until clear. After removal of the supernatant, beads were washed twice with fresh 80% ethanol and elution performed in 13 µL IDTE (IDT). The indexing PCR reaction was set up with 5 µL 4X rhAmpSeq Library Mix 2 (IDT), 4 µL i5/i7 Illumina sequencing indexes at 5 µM each, and 11 µL cleaned-up PCR1 product and was carried out as follows: 95°C for 3 min, 15 cycles of (95°C for 15 s, 60°C for 30 s, 72°C for 30 s), and finally 72°C for 1 min and hold at 4°C. PCR products were cleaned-up using 1 × Ampure XP beads (Beckman Coulter) as described above, using 20 µL room temperature beads and an elution volume of 22 µL IDTE. Quality control of the libraries was performed using the Qubit (ThermoFisher Scientific) and D1000 Tapestation (Agilent). Equal ng aliquots of each library were then pooled together and sequenced using the appropriate Illumina sequencer, aiming to achieve roughly 70,000 reads per amplicon sequenced. Seven of the 28 off-target sites failed to have >10,000 reads and were excluded from the analysis (Supplemental Table 2).

Sequencing reads were first pre-processed to trim low-quality base calls. Subsequently, paired-end reads were stitched to create consensus reads with adjusted base-quality scores, and successfully stitched reads were aligned to the human reference genome. Base call frequencies corresponding to each position in all on-target and candidate off-target sites were calculated from the read alignments. Base call frequencies for treated and untreated samples from the same donor were compared and an odds ratio quantifying the enrichment of each observed variant in the treated sample was calculated. A Fisher’s exact test was used to assess statistical significance.

### Tissue preparation for histology

All samples were fixed in 4% paraformaldehyde (PFA) in PBS. Human retinal and RPE/choroid explants (Figure 2D and F and Supplemental Figures 10A,C and 11A,C) were fixed for 30 min at room temperature. Mouse eye cups (Figure 3B) were fixed for 1 h at room temperature. Human retinal organoids (Supplemental Figure 4A, 10B and 11B) were fixed for 4 h at 4°C. The edge of the NHP bleb (Figure 4B) was fixed overnight at 4°C. After fixation, samples were washed with PBS and cryoprotected in 30% sucrose overnight at 4°C. Samples were then stored at −80°C until processing.

### Tissue embedding and cryosectioning

Human retinal organoids and NHP retinas were embedded in 7.5% gelatin and 10% sucrose in PBS. Mouse eye cups were embedded in a 1:1 mixture of 30% sucrose and O.C.T. media (VWR). Tissues were cryosectioned into 20-25 µm thick sections using a MICROM International cryostat (Walldorf, Germany) and mounted onto Superfrost Plus slides (Thermo Fisher Scientific). Sections were dried overnight at room temperature and stored at −80°C until use.

### Immunofluorescence staining of tissue cryosections

Slides were dried for 1 h at room temperature and then rehydrated in PBS. Samples were blocked in a ‘blocking buffer A’ (PBS supplemented with 10% normal donkey serum, 1% (w/v) bovine serum albumin, 0.5% Triton X-100 in PBS, and 0.02% sodium azide (all from Sigma-Aldrich)) at room temperature for 1 h. Primary antibodies were diluted in 200 µL of ‘blocking buffer B’ (PBS supplemented with 3% normal donkey serum, 1% BSA, 0.5% Triton X-100 in PBS and 0.02% sodium azide) and samples incubated at room temperature in a wet chamber overnight. Slides were washed 3 x 15 min in PBS with 0.05% Triton X-100. Secondary antibodies were diluted in 200 µL of ‘blocking buffer B’ and samples incubated for 2 h at room temperature in the dark. Slides were washed 2 x 10 min in PBS with 0.05% Triton X-100 and 1 x 10 min in PBS and were coverslipped with ProLong Gold (Thermo Fisher Scientific).

### Immunofluorescence staining of tissue cryosections with heat-induced antigen retrieval

To prevent tissue sections from falling off the slides during the antigen retrieval, slides were baked at 60°C for 1 h and postfixed for 15 min on ice with 4% PFA in PBS. Samples were washed shortly in PBS and permeabilized for 15 min with 0.5% Triton X-100 in PBS. The slides were then transferred to a plastic container filled with preheated 1X antigen retrieval buffer (100X Tris-EDTA Buffer; pH 9.0) in water and kept in a steamer for 20 min. After washing in PBS, the samples were stained as described above.

### Immunofluorescence staining of whole-mount human retinas

Whole-mount samples were freeze-thawed three times on dry-ice and washed in 500 µL PBS for 15 min in a 24-well plate. Samples were then blocked in 200 µL ‘blocking buffer A’ for 1 h on a shaker at room temperature in the dark. Primary antibodies were diluted in 200-250 µL of ‘blocking buffer B’ and samples incubated for 3-4 days on a shaker at room temperature in the dark. They were then washed three times for 10-15 min each in 500 µL PBS and incubated with 200 µL of secondary antibodies diluted in ‘blocking buffer B’ for 1.5 h on a shaker at room temperature in the dark. After three washes in 500 µL PBS for 10-15 min each, samples were mounted on slides with ProLong Gold (Thermo Fisher Scientific).

### Immunofluorescence staining of whole-mount samples with heat-induced antigen retrieval

To prevent whole-mount samples from folding during the antigen retrieval, human or *Abca4^ms1961G/G^* wild-type mouse retinas were placed into a hybridization chamber (Grace Bio-Labs) and permeabilized inside the chamber for 15 min with 0.5% Triton X-100 in PBS. After washing with PBS, the chamber was filled with 1X antigen retrieval buffer in water (100X

Tris-EDTA Buffer; pH 9.0), sealed and transferred to a plastic container that was then placed into a steamer for 20 min. The whole mounts were finally washed in PBS and stained as described in the section above.

### RNAScope

Cultured human retina samples were sectioned by cryosectioning. A Hs-ABCA4 probe (ACD Bio) was used to label *ABCA4* mRNA molecules, and the reaction was developed using the RNAScope 2.5 HD Reagent Kit according to the manufacturer’s instructions. Secondary staining was carried out with the Opal 690 dye at a 1:5,000 dilution in TSA buffer. Images were captured with an Olympus LSM710 laser scanning confocal microscope.

### Targeting efficiency of cone and rod photoreceptor cells by AAV capsid

We analyzed six to twelve individual regions of interest (ROIs) per retinal whole mount (*n* = 2 (AAV5) and *n* = 3 (AAV9-PHP.eB) human retinal explants) (Figure 2F). Quantification was performed using the ImageJ Plugin ‘Cell counter’ on two separate z-plane images. A cone-rich layer (layer 1) was extracted based on arrestin3 expression and a rod-predominant layer (layer 2) was extracted based on the absence of arrestin3 expression and the presence of a rhodopsin signal. Cone targeting efficiency was quantified from layer 1 by determining the percentage of eGFP positive cells in the overall arrestin3-positive cone population. Rod targeting efficiency was quantified from layer 2 by determining the percentage of eGFP positive cells in the overall Hoechst-positive rod population.

### Co-expression efficiency of AAV9-PHP.eB-SABE1(N) and AAV9-PHP.eB-SABE1(C) in mouse photoreceptor cells

We analyzed six regions of interest (ROIs) per mouse-retinal and RPE/choroid/sclera whole mounts (*n* = 1). Quantification was performed with an ImageJ Plugin “Cell counter” in a single z-plane image from the outer nuclear layer or the RPE layer. Co-expression efficiency was quantified by determining the percentage of cells co-expressing ABE(N) and ABE(C) in the overall Hoechst-positive population.

### Statistical analysis

We used the following statistical tests: Linear (mixed) models including 3-way mixed-effect ANOVA and 2-way mixed-effect ANCOVA with time as continuous covariate, or t-test (1-way ANOVA). The results were corrected for multiple testing either by Dunnett’s- or Tukey’s correction. A random effect has been introduced into the models when A7 editing and A8 editing have been applied to the same samples, to correct for the correlation of the resulting outcomes. In these cases, the estimates of the correlation between the outcomes of A7 editing and A8 editing on the same samples are reported in Supplemental Table 1. All tests were performed with the statistical software package R (version 4.2.2). *P*-values are stated in Supplemental Table 1.

## AUTHOR CONTRIBUTIONS

A.M. designed and conducted experiments, analyzed, interpreted data and wrote the paper. J.S., W.S., M.W., C.P-W., D.B. designed experiments, analyzed and interpreted data. B.K., J.M., S.H. P.G., M.D., P.B. A.K., S.P., A.G., L.J-K., A.K. performed experiments and analyzed data. T.V. provided AAV vector production and analyzed data. L.B., Q.X, M.Q., C.R., and C.S.C. analyzed data. D.P.M., F.K. A.S. performed human retina experiments. Z.Z.N. provided human retina. P.H. and T.A. performed NHP injections and interpreted data. P-H. M. and L.F. helped to design NHP experiments and interpreted data. M.C. provided statistical analysis. M.R., H.P.N.S., G.C. interpreted data. B.R. analyzed data, interpreted data and wrote the paper. B.G. designed experiments, interpreted data and wrote the paper.

## DATA AND MATERIALS AVAILABILITY

All data are available in the manuscript or the supplementary materials.

